# Mutations change excitability and the probability of re-entry in a computational model of cardiac myocytes in the sleeve of the pulmonary vein

**DOI:** 10.1101/2021.09.24.461636

**Authors:** Karoline Horgmo Jæger, Andrew G. Edwards, Wayne R. Giles, Aslak Tveito

## Abstract

Atrial fibrillation (AF) is a common health problem with substantial individual and societal costs. The origin of AF has been debated for more than a century, and the precise, biophysical mechanisms that are responsible for the initiation and maintenance of the chaotic electrochemical waves that define AF, remains unclear. It is well accepted that the outlet of the pulmonary veins is the primary anatomical site of AF initiation, and that electrical isolation of these regions remains the most effective treatment for AF. Furthermore, it is well known that certain ion channel or transporter mutations can significantly increase the likelihood of AF. Here, we present a computational model capable of characterizing functionally important features of the microanatomical and electrophysiological substrate that represents the transition from the pulmonary veins (PV) to the left atrium (LA) of the human heart. This model is based on a finite element representation of every myocyte in a segment of this (PV/LA) region. Thus, it allows for investigation a mix of typical PV and LA myocytes. We use the model to investigate the likelihood of ectopic beats and re-entrant waves in a cylindrical geometry representing the transition from PV to LA. In particular, we investigate and illustrate how six different AF-associated mutations can alter the probability of ectopic beats and re-entry in this region.

## 1 Introduction

Atrial fibrillation (AF) is the most common arrhythmia in adult humans [1, 2]. Several ion channel mutations, along with a range of other genetic variants and broader risk factors, are known to increase the likelihood of developing AF [3, 4, 5]. The lifetime risk of developing AF after the age of 55 is astonishingly high (37 %, see [6]). Treatment of AF remains a significant challenge [7, 8], perhaps because the biophysical processes initiating and maintaining AF remain unclear [1]. Numerical computations have aided in developing our present understanding of the atrial action potential both for a single atrial myocyte [9, 10, 11] and in tissue [12, 13, 14, 15, 16, 17].

It is well established that AF often is initiated in the pulmonary vein ‘sleeves’ of the left atrium [18, 19, 20], where thin layers of myocardium merge with the fibrous tissue of the pulmonary veins. The densities of certain ion channels expressed in the sarcolemma of myocytes in this region are distinctly different from the rest of the atria [21, 22, 23, 24] and the myocyte-to-myocyte electrical coupling through gap-junctions can vary significantly [12, 20, 25]. The mathematical approaches commonly used for simulating electrical activity in atrial tissue are the bidomain or monodomain models [26, 12, 14, 27]. However, both of these models require spatial averaging over hundreds of cells and it is therefore not possible to model the significant cell-to-cell variations in the electrophysiological properties that are characteristic of the cardiac myocytes located in the ‘sleeve’ of tissue at the transition from the pulmonary vein to the left atria. In order to allow for cell-to-cell variation in ion channel densities and assess the effects of random variation in gap-junction coupling between the myocytes, we have applied a recently developed mathematical model [28, 29, 30, 31, 32] that represents each my-ocyte individually. This is referred to as the EMI model since it explicitly represents the extracellular (E) space, the membrane (M) and the intracellular (I) space, and thus follows the modeling tradition represented by, e.g., [25, 33]. The average mesh length applied in the EMI model is Δ*x* = 10 *μ*m whereas Δ*x* = 0.25 mm is the standard mesh length for the bidomain model; see e.g. [34, 35, 36, 37, 32, 28]. Consequently, the size of one mesh block is about 1 pL for the EMI model and 15600 pL for the bidomain model. This should be compared to 16 pL which is the volume of an atrial myocyte used in the computations below. This resolution allows EMI to interrogate physiological properties at another scale than is possible with the Bidomain model, but at the cost of a 15600-fold increase in the number of mesh blocks. Note that since convergence of the bidomain model is achieved at the mesh size ~Δ*x* = 0.25 mm, further physiological details of cannot be achieved by decreasing the mesh parameter using this model.

A range of genetic variants and mutations are known to increase the probability of developing AF, and a number of these occur in genes encoding the major cardiac ion channels [5, 3, 4]. In some cases, the electrophysiologic outcomes of these mutations have been well characterized, and provide a basis for asking whether a computational framework can be used to identify and characterize clinically meaningful arrhythmogenesis in the pulmonary vein sleeve. To implement such a framework using the EMI model, adequate data sets are available for the following mutations: N588K [38, 39, 40], A545P [41], E229V [42], E375X [43], A130V [44], c.932delC [45]. Here, we investigate how each of these mutations change the likelihood of initiating or maintaining reentry in a model of the specialized myocytes located at the transition from the pulmonary vein (PV) to the left atrial myocardium (LA). Furthermore, we examine how each mutation can alter the excitability of quiescent tissue in this region, and thus change the probability of triggering ectopic beats. In this numerical model, we introduce electrophysiologic heterogeneity in two ways: (1) by allowing each myocyte to be defined as a random combination of previously measured electrophysiologic phenotypes of PV and LA myocytes, and (2) by allowing electrical coupling to vary randomly among adjacent myocytes.

Our computations reveal that the effect of the selected mutations on the biomarkers of the action potential and cytosolic [Ca^2+^] varies significantly. None of the mutations markedly alter the resting membrane potential (RMP), whereas most have significant effects on the maximal velocity of the action potential (AP) upstroke, and AP duration (APD). As expected, we observed re-entry for a range of mutations that either slowed conduction velocity or shortened APD, namely N588K, E299V, A130V and c.932delC. Re-entry was not observed for A545P or E375X (and not for the baseline WT model). Tissue excitability was unchanged for most mutations, but significantly decreased for A130V, and significantly increased for c.932delC. Together, these results suggest that this high resolution (EMI) modeling approach can be applied to simulating the heterogeneous PV-LA junction, and thereby identify and discriminate the arrhythmogenic influence of clinically meaningful perturbations to myocyte electrophysiology.

## 2 Methods

In this section, we describe the models used in our simulations of the PV sleeve. First, we describe membrane models used to account for the ionic currents and intracellular ion concentrations of LA and PV cardiomyocytes. Next, we explain how each of the six considered mutations are represented in the models and the setup used for the finite element simulations of the EMI model. Finally, we provide the definitions of the biomarkers used in our computations.

### 2.1 Membrane models for PV and LA cardiomyocytes

To represent the resting membrane potential and action potential of both human LA and PV cardiomyocytes (CMs), we have utilized adjusted versions of the previously published base model [46, 39]. Versions of our model have previously been used to simulate human induced pluripotent stem cell-derived CMs (hiPSC-CMs) and healthy adult human, canine, rabbit, guinea pig and zebrafish ventricular CMs [46, 47]. In addition, this formalism has been used to study the short QT syndrome mutation N588K in ventricular CMs [39, 48]. The base model used in this study is similar to the version used in [39, 48]. Additional membrane currents (e.g., *I*_Kur_) have been added to more accurately represent the AP waveform and underlying dynamics of atrial myocytes. The full base model formulation used here is found in the Supplementary Information. Unless otherwise specified, simulations characterizing the behavior of this model were performed at 1 Hz pacing frequency.

In order to represent the important differences between LA and PV myocytes, we have used published experimental data directly assessing differences in the membrane currents in myocytes isolated from both regions in the adult canine heart [22]. The only differences required to capture these effects in the LA and PV versions of the model are the maximum conductances of five transmembrane currents. Each has been shown to differ between LA and PV cells in [22]. The factors used to scale the conductance of the currents in the PV version of the model from the conductance of the currents in the LA version of the model are given in Table 1.

**Table 1:** Scaling factors for the currents of the pulmonary vein (PV) version of the base model, taken from [22].

| Parameter | PV scaling factor |
| --- | --- |
| $g_{K1}$ | 0.58 |
| $g_{Kr}$ | 1.5 |
| $g_{Ks}$ | 1.6 |
| $g_{to}$ | 0.75 |
| $g_{CaL}$ | 0.70 |

### 2.2 Modeling the AF-associated mutations

In this study, we have considered six different mutations that have been linked to AF [3]. Five of these mutations alter the function of specific ion channels in the sarcolemmal membrane of CMs and one affects the function of gap junction elements (connexins) that couple neighbouring CMs. We assume the mutations affect the function of individual channel proteins, and that the function of these proteins and functional effects of the mutations do not differ between LA and PV myocytes. Therefore, the effect of the mutations is modeled in the exactly same manner for the LA and PV cases. Below, we list the selected mutations and explain how each is represented in the models. This information is summarized in Table 2. For all the cases, the models for the wild type (WT) version of the channels are unchanged.

**Table 2:** Summary of how each of the selected mutations that have been linked to atrial fibrillation are represented in the LA and PV models. Here, GOF refers to gain-of-function mutations, and LOF refers to loss-of-function mutations. Complete mathematical specifications of each of these modified currents are given in the Supplementary Information.

| Mutation | Effect | Modeling | Ref. |
| --- | --- | --- | --- |
| N588K | $I_{Kr}$ GOF | Transition rates between the open and inactivated states adjusted | [38, 40] |
| A545P | $I_{to}$ GOF | $g_{to}$ increased by 75%<br>$\tau_{r_{to}}$ increased by 15% | [41] |
| E299V | $I_{K1}$ GOF | $I_{K1}$ reformulated | [42] |
| E375X | $I_{Kur}$ LOF | $g_{Kur}$ reduced by 90% | [43] |
| A130V | $I_{Na}$ LOF | $g_{Na}$ reduced by 75% | [44] |
| c.932delC | $I_{gap}$ LOF | $G_{gap}$ reduced by 75% | [45] |

#### 2.2.1 N588K

The N588K gain-of-function mutation in the KCNH2 gene encoding the ion channels responsible for the rapidly activating delayed rectifier K^+^ current, *I*_Kr_, is associated with short QT syndrome type 1 and other cardiac arrhythmias, including atrial fibrillation [49, 40]. McPate at al. [38], have published detailed measurements of the difference between WT and N588K *I*_Kr_. We have fitted the *I*_Kr_ Markov model formulation from [50] to these measurements. Specifically, the N588K mutation is represented by altering the transition rates between the inactivated and open states of the Markov model. The full formulation of the WT and N588K versions of the *I*_Kr_ model are specified in the Supplementary Information.

#### 2.2.2 A545P

The A545P gain-of-function mutation occurs in the KCND3 gene encoding the Kv4.3 alpha subunit of channels carrying the transient outward K^+^ current *I*_to_, and is associated with AF [41]. A545P mutant channels exhibit increased peak current, and slowed current inactivation. In accordance with measurements published by Olesen et al.[41], we represent the mutation by increasing the maximum *I*_to_ conductance by 75% and the time constant of inactivation by 15%.

#### 2.2.3 E299V

The E299V gain-of-function mutation in the KCNJ2 gene impacts ion channels responsible for the time-independent inwardly rectifying K^+^ current, *I*_K1_. This mutation is associated with short QT syndrome type 3 and increased incidence of arrhythmias, including AF [42]. To represent the mutation in the models, we use the WT and E299V versions of the atrial *I*_K1_ formulations provided in [42]. These formulations are based on the Grandi et al. human atrial myocyte model [51] and are fitted to WT and E299V *I*_K1_ measurements from [42]. The full formulation of the WT and E299V versions of the *I*_K1_ model are specified in the Supplementary Information.

#### 2.2.4 E375X

The E375X mutation in the KCNA5 gene exerts loss-of-function effects in the Kv1.5 ion channels responsible for much of the measurable ultra-rapidly activating delayed rectifier K^+^ current, *I*_Kur_, and has been causally linked to idiopathic lone AF [43]. Based on measurements of WT and E375X *I*_Kur_ from [43], we represent this mutation in our model by reducing the maximum conductance of *I*_Kur_ to 10% of the WT value.

#### 2.2.5 A130V

A130V is a loss-of-function mutation in SCN3B encoding the *β*3 subunit of the Na^+^ channel complex that is responsible for *I*_Na_, and was uniquely identified in a patient with lone AF from a study of 477 AF patients of Han Chinese descent [44]. Based on measurements of WT and A130V *I*_Na_ from [44], we represent this mutation in the model by reducing the maximum conductance of *I*_Na_ to 25% of the WT value.

#### 2.2.6 c.932delC

The c.932delC mutation in the GJA1 gene encoding the gap junction protein connexin 43 (Cx43) has been identified as a genetic mosaicism underlying lone AF in a patient for whom cardiomyocyte-specific expression of this variant was responsible for the pathology [45]. Based on measurement of the gap junction conductance for WT and the c.932delC mutation from [45], we represent this mutation in the associated model by reducing the gap junction conductance (*G*_gap_, see (2), below) to 25% of the WT value.

### 2.3 Finite element model of the PV ‘sleeve’

We perform finite element simulations of a defined portion of the ‘sleeve’ of the PV using a model, referred to as the EMI model, which represents the geometry and electric potentials of the extracellular space (E), the cell membrane (M) and the intracellular space (I) (see, e.g., [31, 52, 53, 28, 29, 32]). This model takes the form

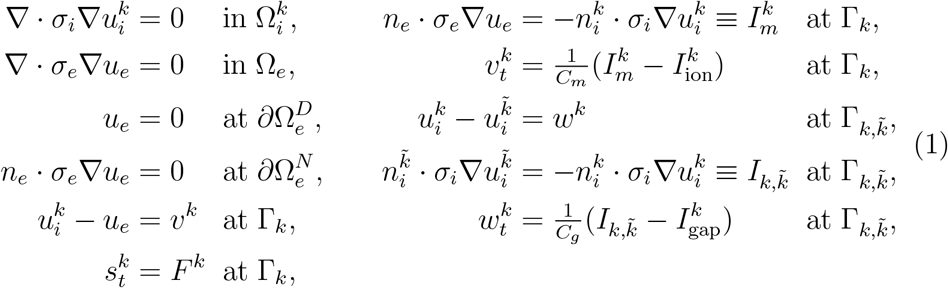

for all myocytes *k* and neighboring myocytes 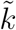 [31]. Here, *u_e_* (in mV) is the electric potential in the extracellular space, Ω_*e*_, and 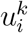 (in mV) is the electric potential in the intracellular space of myocyte *k*, 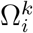. Moreover, 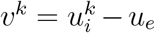 (in mV) is the transmembrane potential of myocyte *k*, defined at the membrane 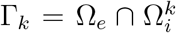, and 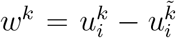 (in mV) is the potential difference between myocyte *k* and its neighboring myocyte 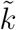, defined at the intercalated disc 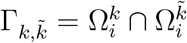. Furthermore, *σ_i_* and *σ_e_* (in mS/cm) is the conductivity of the extracellular and intracellular spaces, respectively, and *C_m_* and *C_g_* (in *μ*F/cm^2^) is the specific capacitance of the cell membrane and the intercalated discs, respectively. In addition, 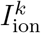 represents the sum of ionic currents through ion channels, pumps and exchangers expressed in the sarcolemma, represented by the base model described in Section 2.1. The state variables of the base model are referred to as *s^k^*, and their dynamics are referred to as *F^k^*. Note that, 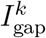 is the current density between neighbouring myocytes, specified by the passive model

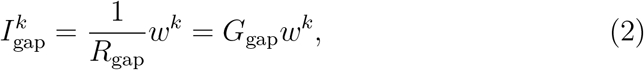

where *G*_gap_ (in mS/cm^2^) is the conductance of the gap junctions and *R*_gap_ (in kΩcm^2^) is the corresponding resistance of the gap junctions. The parameters used in the EMI model simulations are specified in Table 3, and the EMI model equations are solved numerically using an MFEM [54, 55] finite element implementation of the operator splitting algorithm introduced in [30, 56].

**Table 3:** Default parameter values used in the EMI model simulations, see, e.g., [30].

| Parameter | Value |
| --- | --- |
| $C_m$ | $1 \mu\text{F}/\text{cm}^2$ |
| $C_g$ | $0.5 \mu\text{F}/\text{cm}^2$ |
| $\sigma_i$ | $4 \text{ mS}/\text{cm}$ |
| $\sigma_e$ | $20 \text{ mS}/\text{cm}$ |
| $R_{\text{gap}}^*$ (longitudinal) | $0.0005 \text{ k}\Omega \text{ cm}^2$ |
| $R_{\text{gap}}^*$ (transverse) | $0.001 \text{ k}\Omega \text{ cm}^2$ |
| $\Delta t$ | $0.001 \text{ ms}$ |
| $M_{\text{it}}, N_{\text{it}}$ | $1$ |

#### 2.3.1 PV sleeve geometry

We represent essential features of the geometry of the PV sleeve by constructing a collection of coupled myocytes that form a cylinder having a diameter of 1.5 cm (similar to the diameter of a human PV [57]). Each myocyte is shaped as a cylinder with a diameter varying from 13 *μ*m at the cell ends to 14 *μ*m at the cell center, and each myocyte is about 120 *μ*m long [58]. In our formulation, the cylinder of myocytes consists of 393 cells placed longitudinally around the cylinder with 10 rows of cells positioned along the cylinder. The computational mesh of a single cell is illustrated in Figure 1A and an associated cylinder of cells is illustrated in Figure 1B.

**Figure 1:**
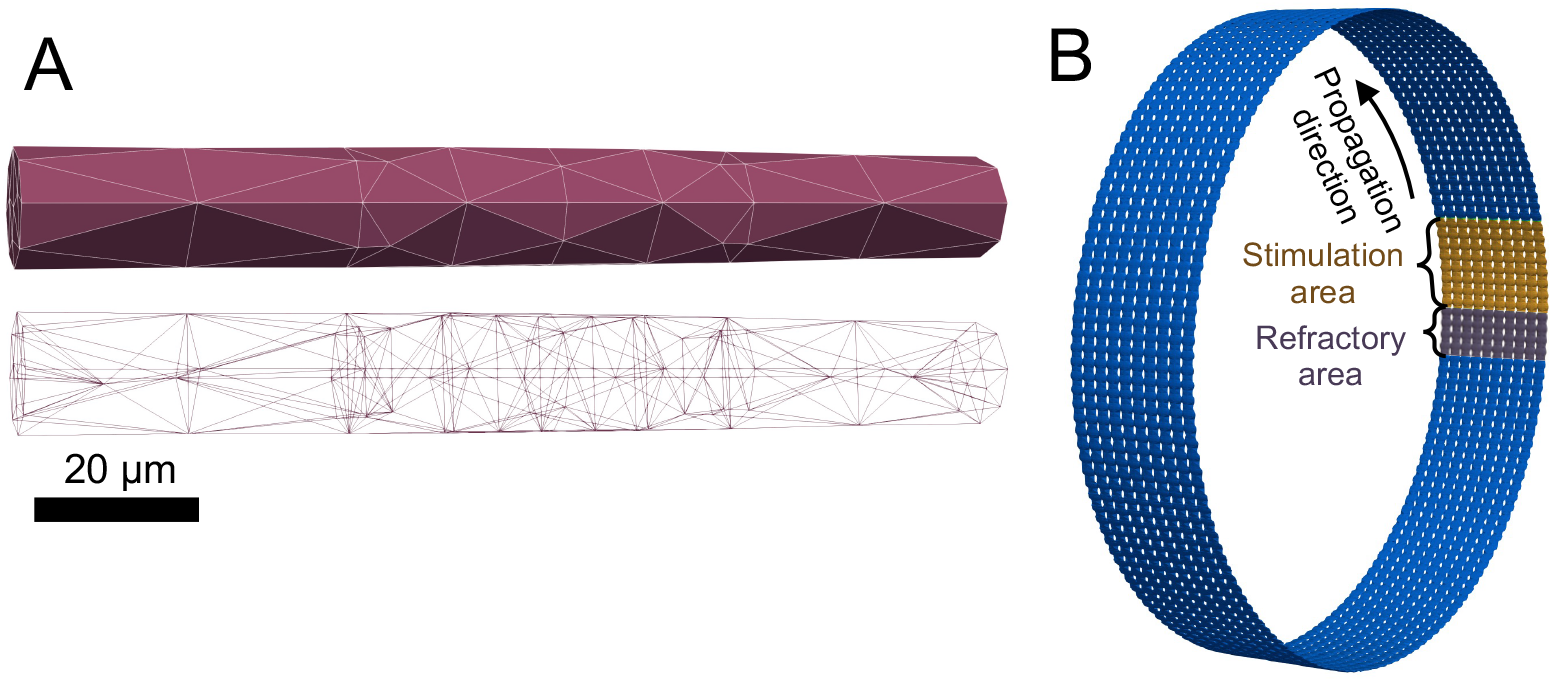
Single myocyte and PV sleeve geometry used in these EMI model simulations. A: The upper illustration shows the surface of a single myocyte within the PV sleeve, and the lower illustration shows the full finite element mesh of the myocyte. B: Illustration of a cylinder of myocytes making up the PV sleeve. Each cell is shaped as illustrated in Panel A. The figure also illustrates the stimulation protocol used in the simulations performed to investigate re-entry (see Section 2.4.2). Note that to improve the visibility of the individual cells of the cylinder, the cylinder in Panel B is not shown to scale.

#### 2.3.2 Distribution of cell properties

We assume that the properties of the CMs forming the sleeve of the PV varies between those of PV and LA CMs but that there is no mean gradient from one end of the cylinder to the other. This arrangement is intended to specifically represent the subsection of the sleeve that is at the border between the vein and remote atrial myocardium. We represent this by drawing a number *α* that is 0 or 1 for each of the 393 × 10 myocytes and letting the conductance of the five currents known to be different between PV and LA CMs (see Table 1) for each cell be given by

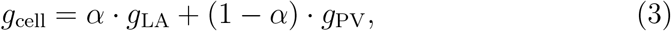

where *g*_LA_ and *g*_PV_ are the LA and PV conductance values, respectively.

Somewhat similarly, the complex localized fibre arrangements and slow and complex conduction that have been observed in the PV sleeve [20] are represented by scaling the gap junction resistance between default values and values corresponding to a considerably reduced cell coupling. This is done by drawing random numbers *β* between 0 and 1 for each intercalated disc and letting

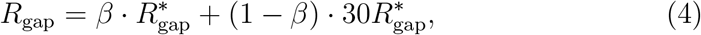

where 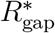 are the default gap junction resistances reported in Table 3.

The numbers *α* and *β* are drawn randomly *once* from a uniform distribution and the same values are reused in all simulations.

### 2.4 Definition of excitability and re-entry

We have performed EMI model simulations based on the PV sleeve to investigate how the selected ion channel or connexin mutations may affect the excitability and/or potential for re-entry in the tissue. Below, we describe the protocols used in these simulations.

#### 2.4.1 Excitability

In order to investigate the excitability of the PV sleeve, we apply a 2 ms rectangular stimulus waveform (*I*_stim_) to 2 × 2 myocytes. We evaluate how intense this depolarizing stimulus current must be to initiate an AP in the tissue, where a larger required *I*_stim_ denotes reduced tissue excitability. We define excitability, *E*, as

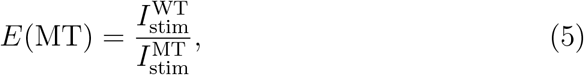

where 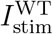 (in A/F) is the required stimulation current in the WT case and 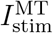 (in A/F) is the required current for the mutation under consideration. Here, *required* refers to the smallest *I*_stim_ capable of initiating a AP.

#### 2.4.2 Re-entry

In order to investigate how each selected mutation affects the likelihood of re-entry, we performed EMI model simulations in which 8 columns of myocytes around the cylinder are stimulated by setting their initial membrane potential to *υ* = −10 mV. In addition, we enforce the condition 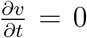 in 4 columns of myocytes on one side of these 8 columns of myocytes, representing a refractory area. This condition is enforced until the propagating AP approaches these myocytes after traveling once around the cylinder. We run the simulation for 500 ms and then determine whether the membrane potential of any of the myocytes is more depolarized than −65 mV anywhere in the cylinder at the end of the simulation. If so, we say that re-entry is obtained in the PV sleeve. The setup used to initiate re-entry is illustrated in Figure 1B.

### 2.5 Defining biomarkers

In the results reported below, we utilize a number of different biomarkers, each computed from the solution of the ordinary differential equations defining the membrane models for LA and PV myocytes and from the EMI model simulations of the PV sleeve with a mix of LA and PV myocytes. In this section, we describe definitions of these biomarkers.

#### I. Electrophysiological

##### a) Resting membrane potential (RMP)

The resting membrane potential (RMP) is defined as the membrane potential obtained 10 ms before the stimulation current that initiates the action potentials is applied.

##### b) Action potential amplitude (APA)

The action potential amplitude (APA) is defined as the difference between the RMP and the maximum value of the membrane potential obtained during an action potential.

##### c) Maximal upstroke velocity (dvdt_max_)

We define the maximal up-stroke velocity (dvdt_max_) as the maximum value of the first derivative of the membrane potential with respect to time during the upstroke of the action potential.

##### d) Action potential duration (APD)

We define the action potential duration at *p* percent repolarization (e.g., APD50, APD80, APD90) as the time between the time when dvdt_max_ is reached and the time during repolarization when membrane potential reaches a value below 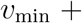 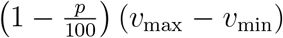, where *υ*_min_ is the minimum value of the membrane potential obtained between two action potentials and *υ*_max_ is the maximum value of the membrane potential during an action potential.

#### II. [Ca^2+^]_*i*_ dynamics

##### a) Ca^2+^ transient amplitude (CaA)

We define the [Ca^2+^] transient amplitude (CaA) as the difference between the maximum value of the cytosolic Ca^2+^ concentration obtained during an action potential and the lowest value obtained between two action potentials.

##### b) [Ca^2+^] transient duration (CaD)

The Ca^2+^ transient duration at *p* percent (e.g., CaD50, CaD90) is defined in the same manner as the APD.

#### III. Conduction velocity (CV)

We compute the conduction velocity in the EMI model simulations used to investigate re-entry by recording the times (after stimulation) at which the myocyte in row 5, column 50 (*t*_1_) and the myocyte in row 5, column 100 (*t*_2_) each first reach positive membrane potentials. The conduction velocity is then computed as

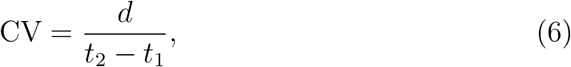

where *d* = 50 · 120 *μ*m = 0.6 cm is the distance between the center of the two myocytes. This corresponds to the longitudinal conduction velocity of the cells, in the direction around the cylinder.

## 3 Results

In this section, we present results from our simulations investigating the effects of selected ion channel protein mutations on the electrophysiological properties of a simulated functional sleeve of the PV. We first present key properties of the wild type models based mainly on the membrane currents of LA and PV myocytes, and we then examine how these properties are affected by each of the six mutations. Next, we investigate the effect of the mutations in EMI model simulations of the multicellular PV sleeve tissue. Specifically, we investigate whether the excitability of the CMs in the sleeve is altered and whether the mutations change the electrophysiological properties of the sleeve in a manner such that a re-entrant wave is sustained.

### 3.1 Properties of membrane models for LA and PV

In order to accurately represent the membrane currents and intracellular Ca^2+^ fluxes of both LA and PV myocytes and relate our findings to previous publications, we use a modified version of the base model from [46, 39]. The differences between the LA and PV versions of our model are closely based on measured differences in transmembrane current densities from [22]. These are specified in Table 1.

Figure 2 shows the AP and cytosolic [Ca^2+^] transient for the LA and PV versions of the base model. In Table 4, we compare a number of biomarkers for the two versions of the model. It is apparent that both base models capture observed semi-quantitative differences between canine LA and PV myocytes from [22]. Specifically, the resting membrane potential is depolarized for the PV model compared to the LA model, and the maximum upstroke velocity of the action potential (AP) is considerably lower for PV than LA. In addition, the APD is longer for LA myocytes than for PV myocytes. All of these characteristics are consistent with the experimental observations [22]. Figure 3 shows the current densities for the different membrane currents of the LA and PV versions of the base model during an action potential.

**Figure 2:**
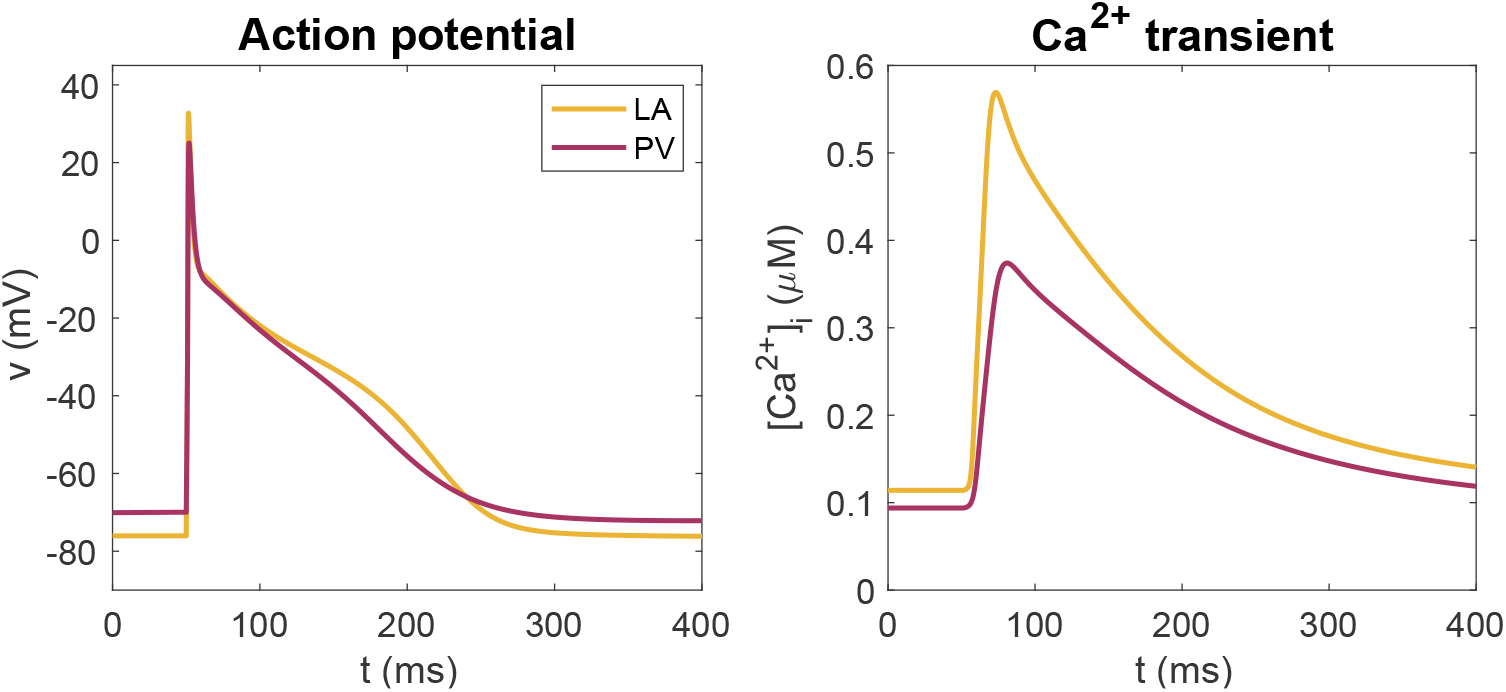
Action potentials (left) and [Ca^2+^] transients (right) for the LA and PV versions of our base model obtained at steady-state in response to a 1 Hz stimulus train.

**Table 4:** Biomarker values for the LA and PV versions of our base model at 1 Hz pacing. Selected biomarkers include the resting membrane potential (RMP), the action potential amplitude (APA), the maximal upstroke velocity of the action potential (dvdt_max_), the action potential durations at 90% and 50% repolarization (APD50 and APD90), the amplitude of the cytosolic Ca^2+^ transient (CaA), and the cytosolic Ca^2+^ transient durations at 90% and 50% (CaD50 and CaD90). These biomarkers are defined in Section 2.5.

|  | LA | PV |
| --- | --- | --- |
| RMP | -76 mV | -70 mV |
| APA | 109 mV | 95 mV |
| $dvdt_{\max}$ | 198 V/s | 101 V/s |
| APD90 | 187 ms | 172 ms |
| APD50 | 48 ms | 51 ms |
| CaA | 0.45 μM | 0.28 μM |
| CaD90 | 277 ms | 323 ms |
| CaD50 | 97 ms | 120 ms |

**Figure 3:**
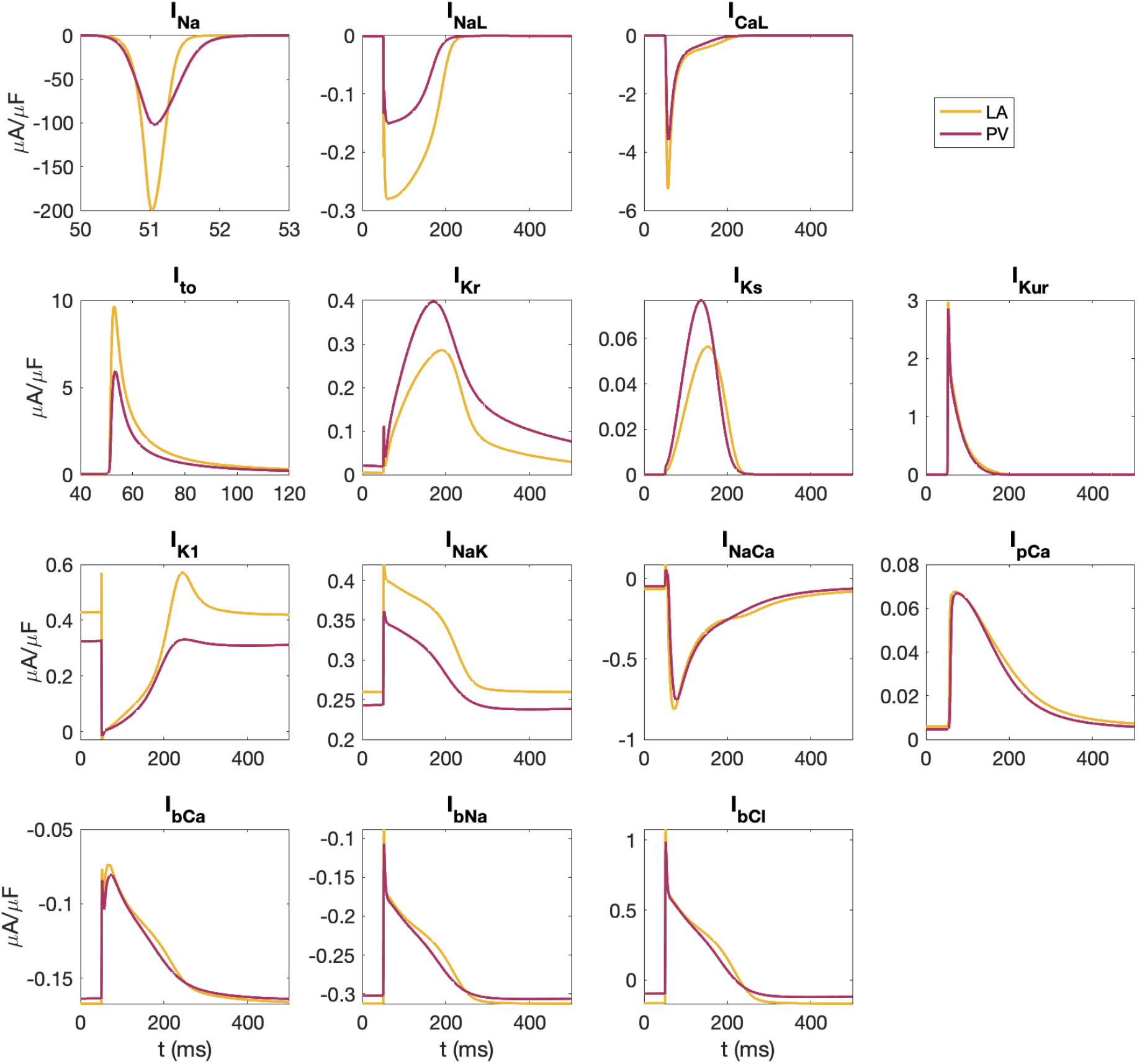
Membrane currents during an action potential in the LA and PV versions of the base model. Definitions and mathematical specifications for each of these currents are provided in the Supplementary Information.

### 3.2 Effect of the mutations on the membrane models

In Figure 4 and Table 5, we investigate the effect of incorporating different mutations in the LA and PV versions of the base model as described in Section 2.2. Note that a number of mutations (N588K, A545P and E299V) lead to a shortened APD for both PV and LA cells, and that the degree of shortening varies among the mutations. For the A130V mutation, we observe a significant decrease in the maximal upstroke velocity. For the E375X mutation, early phase II repolarization is markedly slowed, as is to be expected for this major loss-of-function *I_Kur_* mutant, although terminal repolarization and the indices of later APD remain relatively unaltered. Since the c.932delC mutation affects only gap junction function, which is only implemented for the tissue simulations, none of the parameters of the membrane model are changed for this mutation.

**Figure 4:**
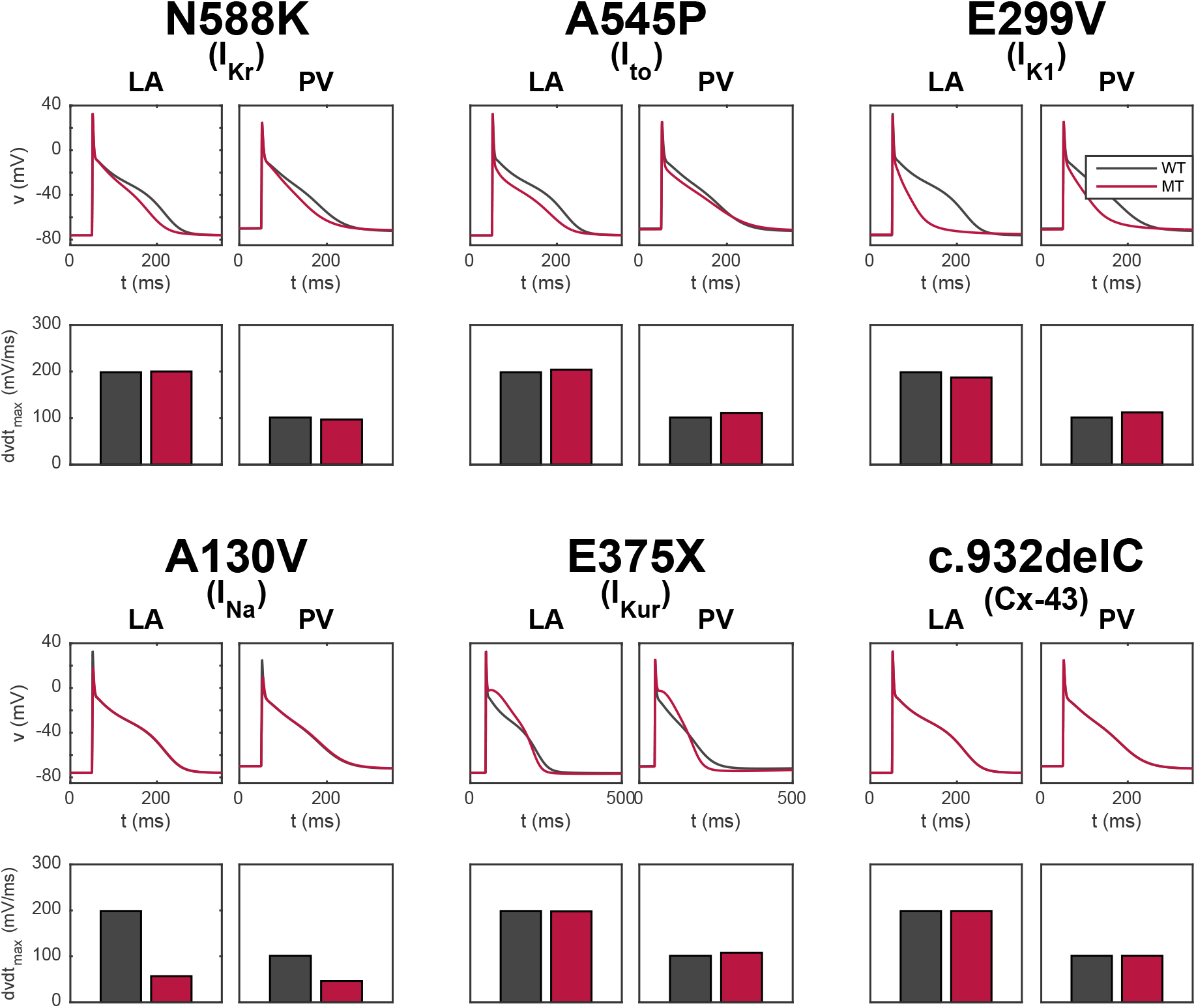
Effects of selected mutations on the AP and maximal upstroke velocity in single myocyte simulations of the LA and PV versions of our membrane model. The black lines and bars refer to WT, and the red lines and bars refer to the mutant cases. Note that the c.932delC mutation (connexin 43) does not affect any of the parameters in the membrane models, but can have a significant effect on intercellular coupling, CV and arrhythmogenic substrate in the multicellular (syncytium) simulations.

**Table 5:**
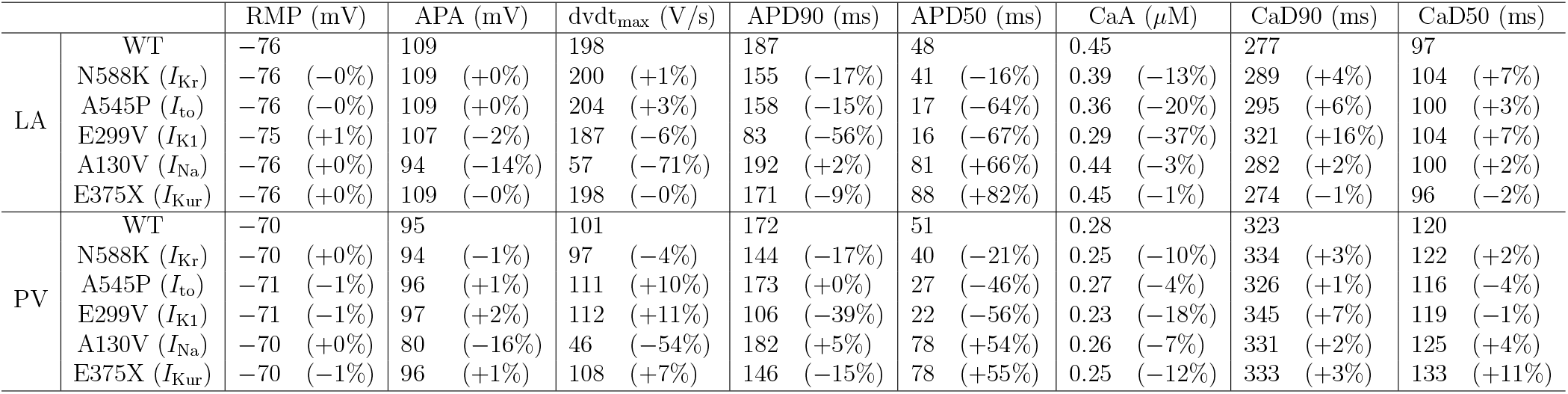
Comparisons between computed WT and mutant biomarker values for simulations of the LA and PV versions of the membrane model driven at 1 Hz. These comparisons are based on changes in the same biomarkers as in Table 4, which are defined in Section 2.5. The numbers in parentheses report the deviation from WT. Note that the c.932delC mutation (in connexin 43) is not included here because these biomarkers are computed from simulations of the single myocyte LA and PV membrane models, and the c.932delC mutation does not affect any of the parameters of the membrane models.

### 3.3 Effect of the mutations on excitability

The effect of each selected mutation on the excitability of the myocytes in the PV sleeve has been investigated using the protocols described in Section 2.4.1. Figures 5 and 6 show the intracellular potential in these simulations at predetermined, fixed points in time. In Figure 5, we observe that a 2 ms stimulus current of 16 A/F is needed to initiate an AP in the WT tissue cylinder, and that this minimum stimulus intensity is unchanged for the N588K, A545P, E299V and E375X mutants. These findings are largely to be expected given that these mutations all impact K^+^ currents, but did not influence resting membrane potential.

**Figure 5:**
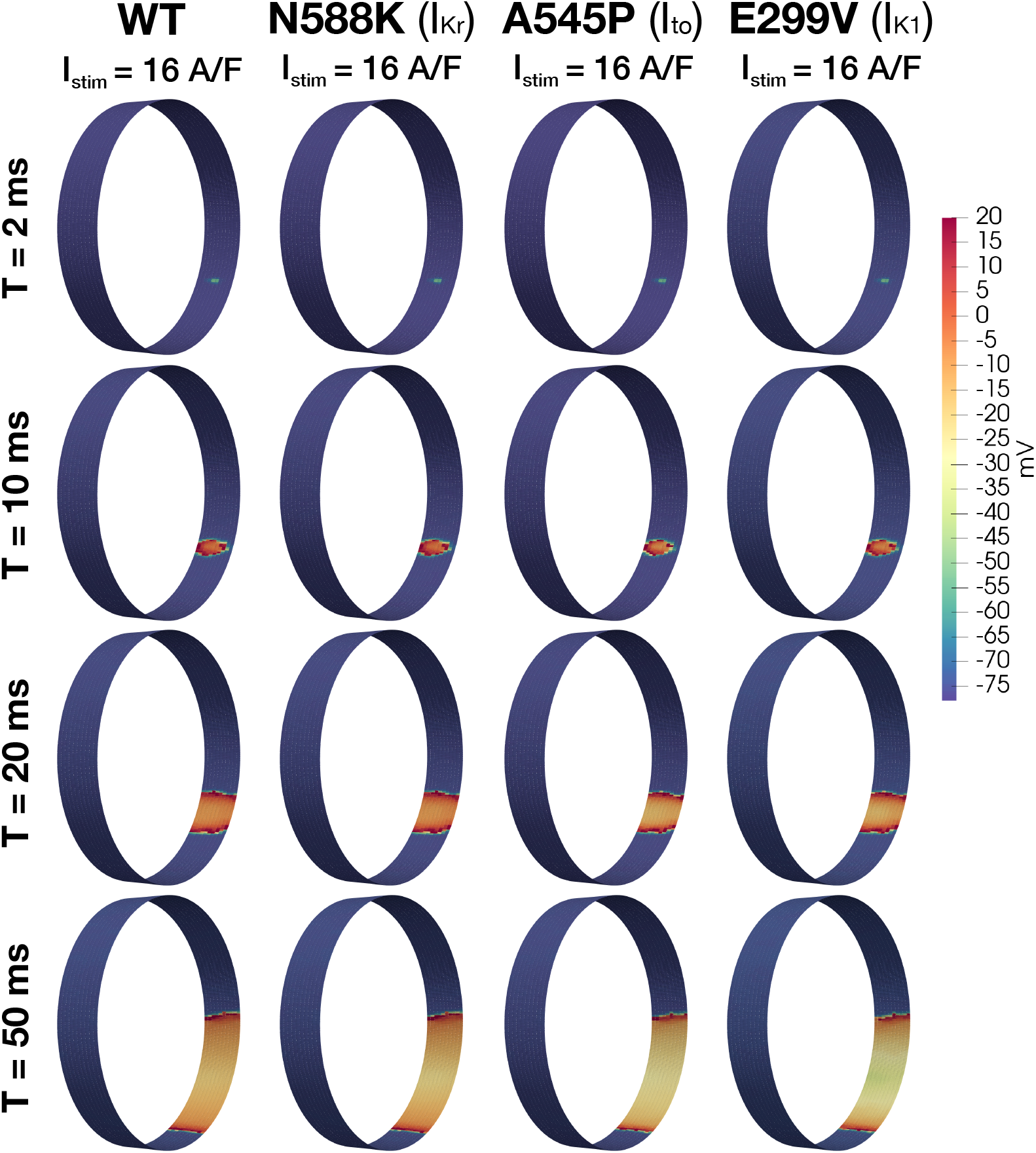
Excitability analysis for WT, and for the N588K, A545P, and E299V mutations. Intracellular potential of myocytes in the PV sleeve is illustrated at four different times (rows) for each mutation (columns). In all cases a regenerative AP was generated and propagated around the tissue cylinder. The stimulus intensity (applied to 2 × 2 cells for 2 ms) required to initiate the propagating action potential in each case is provided in the title for each mutation. Note that to improve the visibility, the longitudinal axis of the cylinder of myocytes has been stretched by a factor of 20.

**Figure 6:**
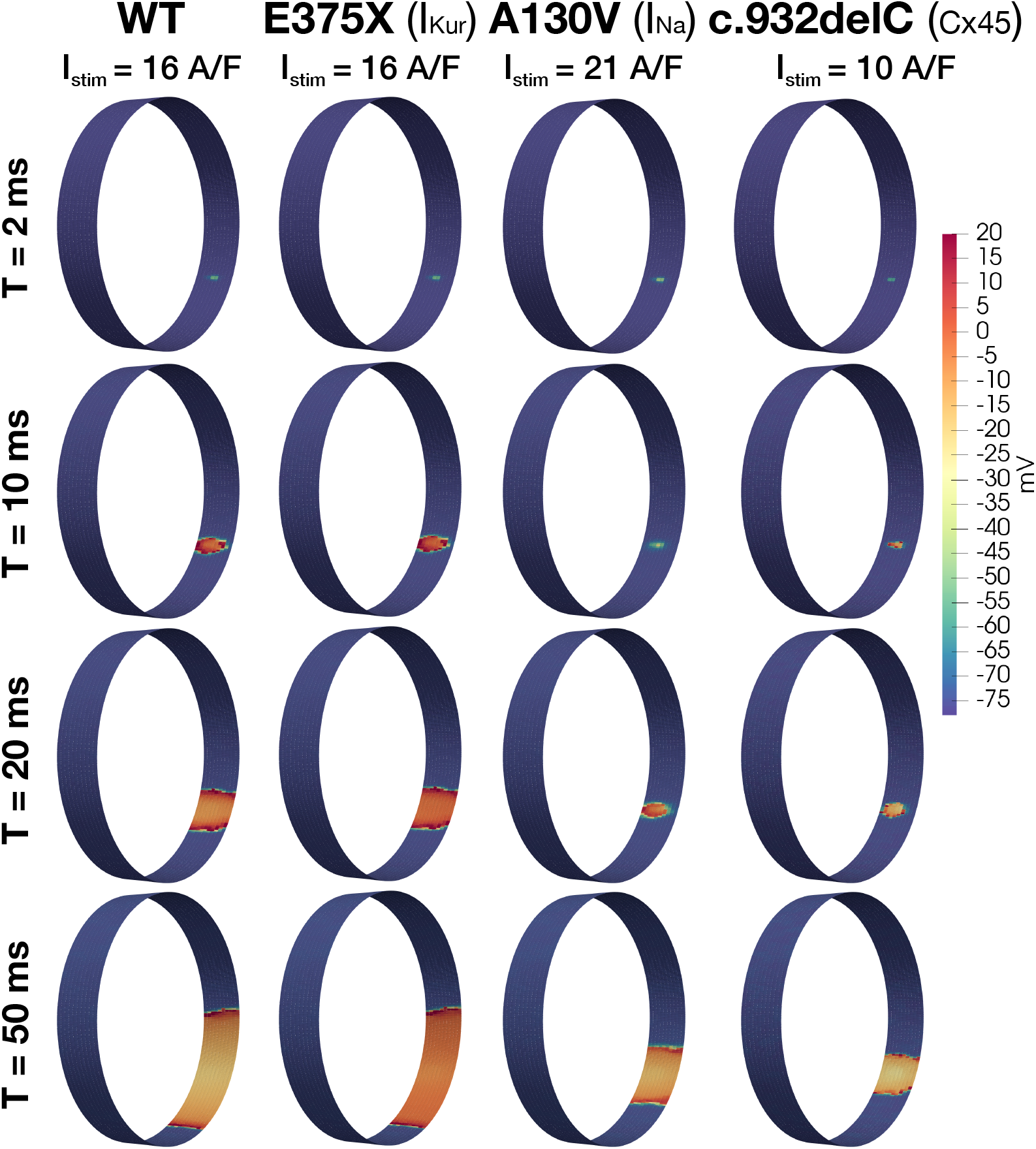
Excitability analysis for WT, and for the E375X, A130V, and c.932delC mutations. This presentation format is the same as that of Figure 5.

In Figure 6, we observe that for the c.932delC mutation, a smaller stimulus current is sufficient to generate a propagating AP, indicating an increased excitability for this mutation. On the other hand, a stronger stimulus is required for the A130V mutation, thus indicating decreased excitability.

### 3.4 Effect of the mutations on re-entry

To investigate how the same mutations affect the initiation and/or maintenance of re-entry, we use the protocols described in Section 2.4.2. Figures 7 and 8 show the intracellular potential of these simulations at some different points in time. In Figure 7, we observe that in the WT case, a propagating wave begins to travel around the tissue cylinder, but upon reaching the stimulation site it encounters a refractory region and terminates. Thus, re-entry is prevented. The same mechanism is observed for the A545P and E375X mutations.

**Figure 7:**
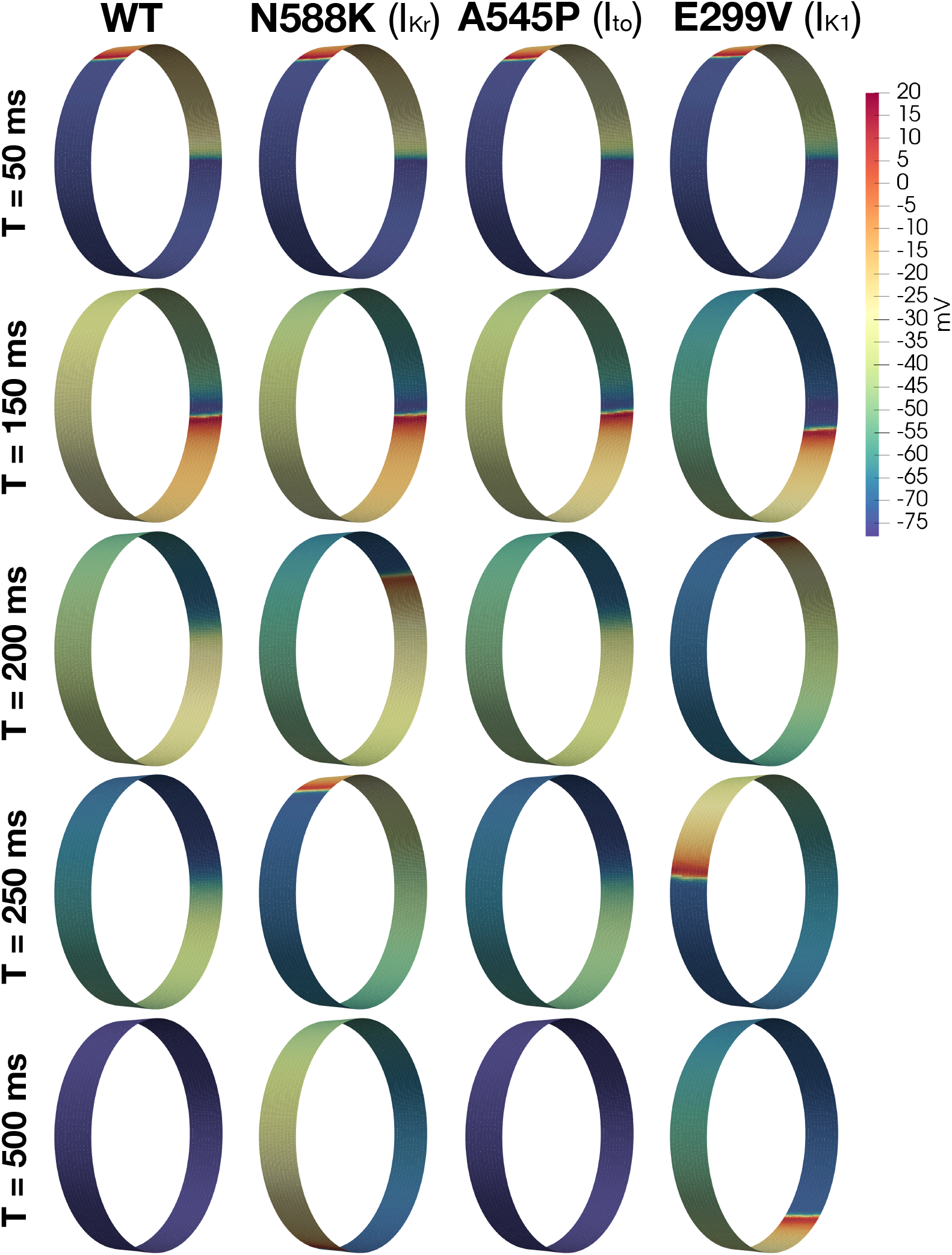
EMI-based multicellular simulations of electrical propagation and re-entry in the cylinder segment of the PV sleeve tissue. We use the simulation protocol described in Section 2.4.2 for investigating re-entry. Data are illustrated for WT, and the N588K, A545P, and E299V mutations (columns). Specifically, we show the intracellular potential of the cells in the PV sleeve at five different times after stimulation (rows). Note that to improve the visual resolution, the longitudinal axis of the PV sleeve cylinder has again been stretched by a factor of 20.

**Figure 8:**
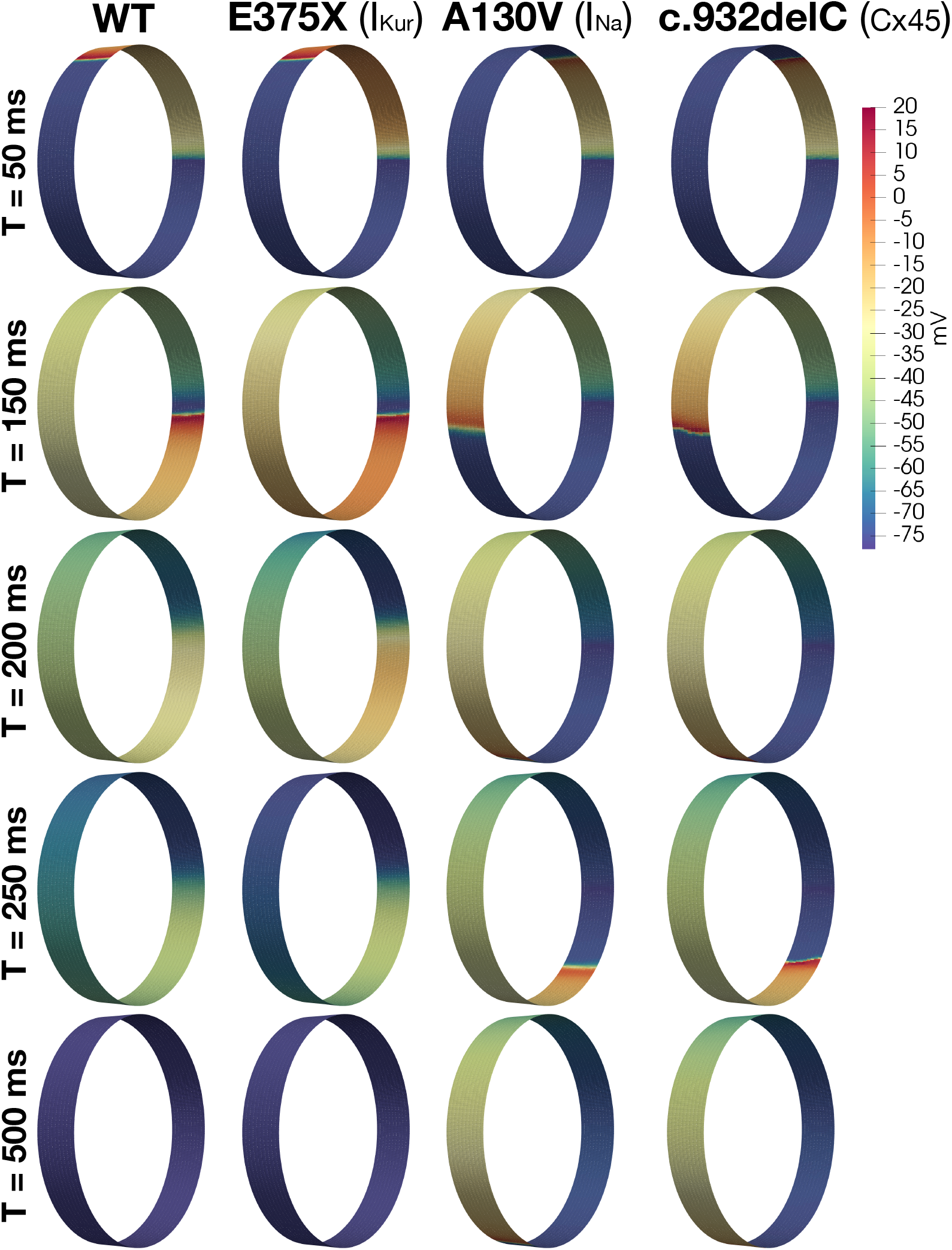
EMI-based multicellular simulations for WT, and for the E375X, A130V, and c.932delC mutations. The format of the figure is analogous to that of Figure 7.

In contrast, the shorter APs and refractory periods for the N588K and E299V mutations (Figure 4), permit the approaching wavefront to re-excite this region. Consequently, the propagating wave continues to travel around the cylinder as a re-entrant wave.

For the A130V and c.932delC mutations, APD is not changed considerably (Figure 4), but for A130V the maximal upstroke velocity is significantly decreased and for the c.932delC mutation, gap junction conductance is reduced by 75%. As a result, CV is considerably slower for both cases, thus permitting the myocytes at the stimulation site to fully repolarize, recover from refractoriness, and re-excite upon arrival of the propagating AP.

### 3.5 Summary of results

The results of these EMI model simulations have revealed important properties and patterns of changes in excitability and re-entry due to AF-associated mutations, and are summarized in Table 6. We observe that for the N588K and E299V mutations, APD (and refractory period) is considerably decreased, and this permits re-entrant behavior. APD is decreased somewhat for the A545P and E375X mutations as well, but not enough to permit reentry. For the A130V and c.932delC mutations the CV is considerably decreased and this predictably allowed re-entrant excitation. In addition, for the A130V mutation, excitability is decreased, whereas for c.932delC excitability is considerably increased.

**Table 6:** Summary of the results of the EMI model simulations of the PV sleeve for WT and eight mutations linked to atrial fibrillation. Definitions of the properties are given in Sections 2.4–2.5, and the numbers in parentheses report the deviation from WT.

|  | APD80 (ms) | CV (cm/s) | Excitability | Reentry |
| --- | --- | --- | --- | --- |
| WT | 162 | 30.5 | 1.00 | N |
| N588K ($I_{Kr}$) | 130 (−20%) | 30.6 (+0%) | 1.00 | Y |
| A545P ($I_{to}$) | 150 (−8%) | 30.6 (+0%) | 1.00 | N |
| E299V ($I_{K1}$) | 78 (−52%) | 29.9 (−2%) | 1.00 | Y |
| E375X ($I_{Kur}$) | 144 (−11%) | 30.5 (−0%) | 1.00 | N |
| A130V ($I_{Na}$) | 176 (+9%) | 15.9 (−48%) | 0.76 | Y |
| c.932delC (Cx43) | 159 (−2%) | 16.2 (−47%) | 1.60 | Y |

## 4 Discussion

The borders between the PV sleeves and working atrial myocardium are small regions of tissue exhibiting pronounced electrophysiologic heterogeneity and structural complexity. Importantly, they are the site of initiation for the majority of spontaneously occurring AF in humans, and electrical isolation of this region remains the single most effective clinical treatment for this remarkably prevalent arrhythmia. Here we have asked whether a recently developed mathematical model (EMI), which is specifically built to interrogate heterogeneities at the level of single cardiac myocytes, is capable of distinguishing clinically meaningful arrhythmogenic influence in this region. We specifically selected a range of genetic mutations known to be the genetic basis of either lone AF, or AF with other comorbid arrhythmias (e.g. short QT syndrome). We then assessed whether the arrhythmogenic impacts of these mutants could be identified by an EMI implementation of a thin cross-section of the PV-LA border. Our model incorporates heterogeneity of myocyte electrophysiology as defined by patch-clamp measurements in isolated PV and LA myocytes, and sharp microelectrode tissue recordings. This approach was readily capable of assessing the proarrhythmic influence of these mutations, and also of revealing one dominant mode of arrhythmogenesis for each. Specifically, it was able to identify and begin to characterize the propensity for re-entrant behavior among mutants that expand the region of excitable tissue (in this small geometry), either by reducing conduction velocity or abbreviating the tissue APD (and hence the tissue refractory period). This computational approach were also able to distinguish the propensity for two mutations impacting *I*_Na_ (A130V) and Cx43 (c.932delC) to markedly alter tissue excitability, albeit in opposite directions. These results suggest that this type of EMI implementation may provide a viable platform for discriminating other clinically important pro- and anti-arrhythmic influences. Most notably for candidate pharmaceutical treatments specific to PV arrhythmogenesis.

### 4.1 The role of high-resolution modeling for understanding arrhythmogenesis in the PV sleeves

Since the initial identification of the PV-LA junctions as the primary sites of AF initiation [18], both clinical and scientific studies have sought to leverage this characteristic to reduce the burden of AF. In basic science investigations, much progress has been made towards understanding the unique characteristics of this region that so strongly predispose it to generating abberant and disruptive electrical activity. Some of these are structural and include the geometry of the tissue (a long, thin cylinder of myocardium directly bordered by the insulating fibrous veinous wall), sharp discontinuities in myocyte fiber direction at the PV-LA border [59, 60], and characteristic fibrous invasions that can create conduction discontinuties and promote local circuit formation [60, 61, 62]. Other important differences are electrophysiologic. PV myocytes have the shortest APs of any myocyte subtype in the human heart [63], and exhibit a clear propensity for triggered activity [64, 65]. Some have suggested that a specific population of myogenic or spontaneously active myocytes exist in the PV sleeves [66], although this has not been broadly confirmed. A range of computational studies have interrogated some of these characteristics and begun to create a biophysically defined heirarchy of arrhythmogenic mechanisms in the PV sleeves. Earlier work from Cherry et al. [67] clearly demonstrated an important interaction of heterogeneity in myocyte-to-myocyte coupling with tissue size for promoting re-entry in a pseudo-PV sleeve. More recent computational efforts have sought to apply clinical imaging data to understand how macroscopic properties of PV electrophysiology and structure interact with electrical activity in the atrial free wall [16, 68, 69, 70, 12, 71, 72]. These approaches often incorporate measured variations in fibre direction, together with electrophysiologic heterogeneities in regions defined at the centimeter scale, in geometries of individual patient atria [70, 16, 71, 12]. Some have also included heuristically defined fibrotic regions within the PV sleeve [12]. Together these studies have suggested a range of structural and dynamic substrate components that are important for permitting macroscopic arrhythmia to initiate (or anchor) around the PV-LA junction, and then propagate into the atrial myocardium. However, because all of these studies employed the bidomain or monodomain models they cannot address electrophysiologic heterogeneities or structural characteristics smaller than a length scale of several millimetres. This precludes investigating a range of properties that rely on local heterogeneities or microscopic fibrotic barriers to create micro-reentrant circuits, or spontaneous foci at the PV-LA border. These proarrhythmic events have been suggested by many studies, and the PV-LA border is among the most probable locations for these mechanisms in humans [60, 61, 62].

To our knowledge, this is the first study that has simulated the PV-LA border at a resolution needed to capture mechanisms that operate at a spatial scale of a single myocyte. Our results provide proof-of-principle that the cell-based EMI framework can discriminate the meaningful arrhythmogenic influences of known AF mutations in simulations of the PV-LA border. We anticipate that the same modeling strategy can be applied to investigate the role of defined local microfibrous structures and realistic heterogeneity in myocyte orientation and coupling with specific electrophysiological characteristics in this critical PV-LA sleeve region.

### 4.2 Simulated pro-arrhythmic properties of AF mutations at the PV-LA junction

While the mutations we have studied here were not specifically chosen for their link to PV slleve arrhythmias, all are channelopathies that have been causally linked to lone AF or AF that is comorbid with other arrhythmia disorders. As such they are electrophysiologically well-characterized and known to predispose to AF without a requirement for other confounding comorbidities. Our implementations of these mutations faithfully recapitulated their electrophysiologic phenotypes at the level of single myocytes. Some of these were quite subtle (A545P, *I*_to_), while others were clearly severe (E299V, *I*_K1_). Our EMI tissue framework was able to distinguish an arrhythmic propensity in 4 of the 6 mutants, and it was only the the E375X (*I*_Kur_) and A545P mutants that did not exhibit clear arrhythmic activity. In all cases this arrhythmic activity was re-entrant, that is, we did not observe spontaneous activity in any of the tissue simulations. However, this likely reflects greater sensitivity of our re-entry protocol, compared to our excitability protocol, for distinguishing arrhythmic potential. In particular, we did not incorporate any instances of the PV or LA myocyte phenotypes that exhibited spontaneous depolarizations or spontaneous APs in the PV-LA tissue. Extending the variability of the myocyte phenotype to permit some instances that are spontaneously active (as would be expected of typical myocyte heterogeneity in tissue) is one direction for the framework that could be of obvious value. We did still observe marked changes in tissue electrotonic properties with our excitability simulations. These suggest that the loss-of-function c.932delC (Cx43) mutant is likely to be particularly predisposed to spontaneous focal activity, whereas *I*_Na_ loss-of-function accompanying A130V may be sufficient to elicit conduction block in certain conditions, although this would require further interrogation. Together, these results suggest that even this simple implementation of the cell-based EMI framework is capable of assessing arrhythmic PV phenotypes involving re-entry, and that more detailed approaches to assessing spontaneous focal activity are likely possible.

### 4.3 Limitations and future directions

While our results suggest that the high-resolution modeling approach permitted by EMI may be very appropriate for interrogating mechanisms of PV arrhythmogenesis, there are areas where we can clearly improve the approach. First, in this study we have not investigated the potential for microfibrotic regions, which are known to occur at the PV-LA border [62], to impact conduction and introduce micro-reentry. This would require significantly larger tissue geometries, which while possible are computationally costly, and thus beyond the initial proof-of-principle we have performed here. Second, while we did implement random heterogeneity that was constrained by the mean properties measured for canine LA and PV myocytes [22], it is likely that real electrophysiologic heterogeneity in the PV region exceeds these constraints. As a cell-based framework, EMI is uniquely well-suited to applying this more realistic heterogeneity, perhaps by invoking populations of myocyte models constructed from the measured variability in those same canine experiments. Third, we have not considered perturbations or abnormalities in calcium homeostasis in these simulations although changes in the intracellular calcium concentration are computed for each myocyte. Such perturbations are well known to play a role in PV-driven arrhythmia [73], and will be a clear priority for future work. Finally, while it would constitute a major extension of the EMI model, it would be highly desirable to implement fully diffusive ionic mass conservation in both the extracellular and intracellular spaces to permit investigation of longer-term changes to ionic homeostasis in these two domains. Major changes in that homeostasis are thought to result from the maintained high frequency firing that is characteristic of AF and to contribute to maintaining AF in patients, and the explicit implementation of the intracellular and extracellular spaces by EMI offers a unique framework for interrogating those mechanisms in a realistic and comprehensive manner.

## 5 Conclusion

This study provides a proof-of-principle demonstration that a cell-based computational modeling framework can be used to distinguish the mechanisms of clinically meaningful arrhythmogenesis in an idealized model of the PV-LA junction.

## 6 Supplementary Information

### 6.1 Base model formulation

In this section, we describe the formulation of the human left atrial and pulmonary vein myocyte versions of our base model. Here, the membrane potential (*υ*) is given in units of mV, and the Ca^2+^ and Na^+^ concentrations are given in units of mM. All currents are given in units of A/F, and the ionic fluxes are expressed as mmol/ms per total cell volume (i.e., in units of mM/ms). Time is given in ms. The parameters of the model are all given in Tables 7–13. In particular, the adjustment factors used to scale the model from the left atrial version to the pulmonary vein version of the model are found in Table 10.

Note that the model formulation is very similar to the base model from [46, 39]. The main difference is that the *I*_Kur_ current is included and that the conductances of the currents and fluxes are adjusted to represent atrial cardiomyocytes.

#### 6.1.1 Membrane potential

In the base model formulation, the membrane potential is governed by

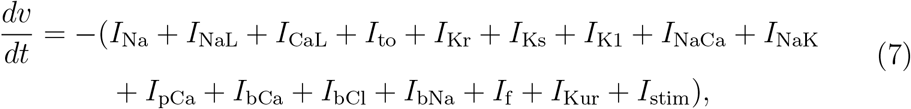

where *I*_Na_, *I*_NaL_, *I*_CaL_, *I*_to_, *I*_Kr_, *I*_Ks_, *I*_K1_, *I*_NaCa_, *I*_NaK_, *I*_pCa_, *I*_bCl_, *I*_bCa_, *I*_bNa_, *I*_f_, and *I*_Kur_ are transmembrane currents that will be specified below and *I*_stim_ is an applied stimulus current. Unless otherwise specified, we let *I*_stim_ be given as a constant current of size −40 A/F applied until the membrane potential reaches a value of −40 mV.

#### 6.1.2 Transmembrane currents

In general, the currents through the voltage-gated ion channels within the myocyte membrane are given on the form

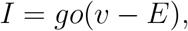

where *g* is the channel conductance, *υ* is the membrane potential and *E* is the equilibrium potential of the channel. Moreover, *o* is the open probability of the channels, which is given on the form *o* = **Π**_*i*_ *z_i_*, where *z_i_* are gating variables. These gating variables are either given as an explicit function of the membrane potential or governed by equations of the form

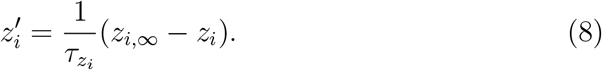

The parameters 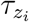 and *z*_*i*,∞_ are specified for each of the gating variables of the model in Table 15.

##### Fast sodium current (*I*_Na_)

The formulation of the fast sodium current
is based on the model formulation given in [74] and is given by

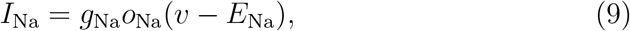

where the open probability is given by

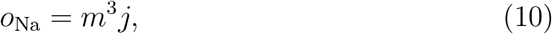

and *m* and *j* are gating variables governed by equations of the form (8).

##### Late sodium current (*I*_NaL_)

The formulation of the late sodium current, *I*_NaL_, is based on [75] and is given by

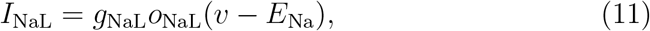

where the open probability is given by

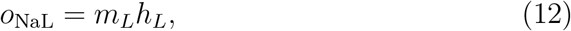

and *m_L_* and *h_L_* are gating variables governed by equations of the form (8).

##### Transient outward potassium current (*I*_to_)

The formulation of the transient outward potassium current, *I*_to_, is based on [76] and is given by

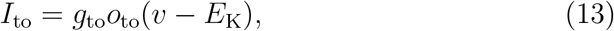

where the open probability is given by

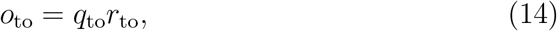

and *q*_to_ and *r*_to_ are gating variables governed by equations of the form (8). For the A545P mutation, *g*_to_ is increased by 75% and 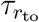 is increased by 15%.

##### Rapidly activating potassium current (*I*_Kr_)

The rapidly activating potassium current, *I*_Kr_, is formulated as a Markov model, based on [50]. The formulation has been fitted to data of WT and N588K *I*_Kr_ currents from [38]. The current is given by

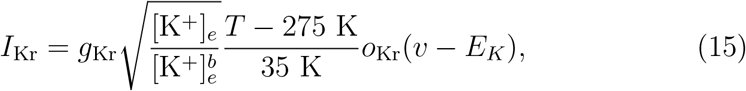

where *o*_Kr_ is modelled by a Markov model of the form

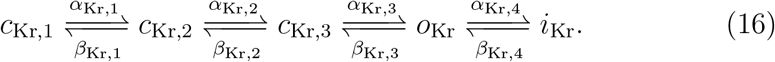

Here, the dynamics of the closed states *c*_Kr,1_, *c*_Kr,2_, and *c*_Kr,3_, the open state *o*_Kr_, and the inactivated state *i*_Kr_ are given by

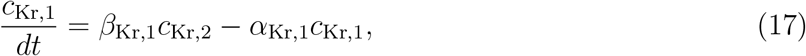

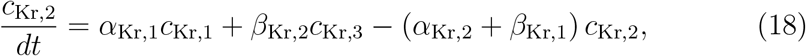

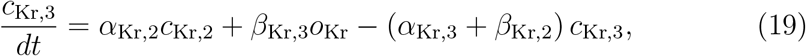

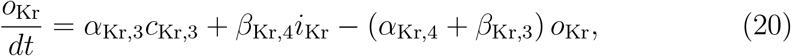

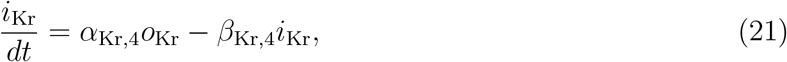

where the transition rates are given by provided in Table 14. The N588K mutation is represented in the model by multiplying *α*_Kr,4_ and *β*_Kr,4_ by 0.25 and 5, respectively.

##### Slowly activating potassium current (*I*_Ks_)

The formulation of the slowly activating potassium current, *I*_Ks_, is based on [74] and is given by

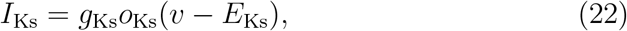

where

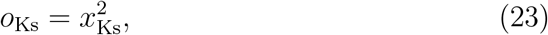

and the dynamics of *x*_Ks_ is governed by an equation of the form (8).

##### Inward rectifier potassium current (*I*_K1_)

The formulation of the inward rectifier potassium current, *I*_K1_, is based on [42, 51], and is given by

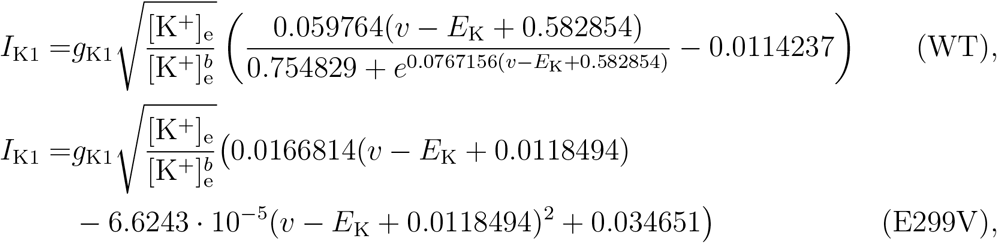

for wild type and the E299V mutation, respectively.

##### Ultrarapid delayed rectifier potassium current (*I*_Kur_)

The formulation of the ultrarapid delayed rectifier potassium current, *I*_Kur_, is based on [77, 78] and is given by

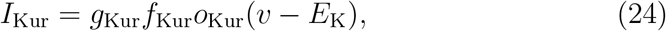

where *f*_Kur_ is given by

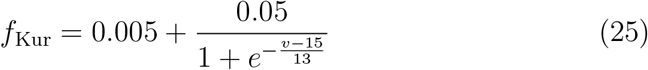

and

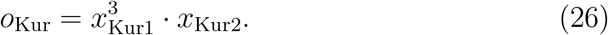

The dynamics of *x*_Kur1_ and *x*_Kur2_ are governed by equations of the form (8). For the E375X mutation *g*_Kur_ is reduced by 90%.

##### Hyperpolarization activated funny current (*I*_f_)

The formulation for the hyperpolarization activated funny current, *I*_f_, is based on [76] and is given by

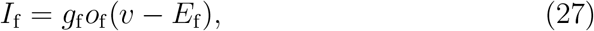

where

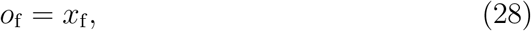

and the dynamics of *x*_f_ is governed by an equation of the form (8).

##### L-type Ca2+ current (*I*_CaL_)

The formualtion for the L-type Ca^2+^ current, *I*_CaL_, is based on the formulation in [74] and is given by

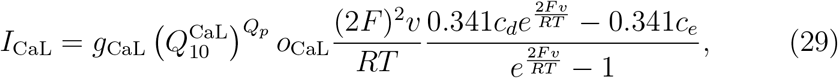

where

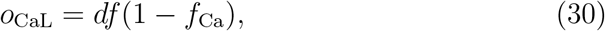

and the dynamics of *d*, *f* and *f*_Ca_ are governed by equations of the form (8). Note that compared to earlier versions of the base model, the dynamics of the *d* gate have now been adjusted in order for the current to behave similarly to other human published atrial membrane models [77, 79, 10].

##### Time-independent background currents (*I*_bCa_, *I*_bBa_, *I*_bCl_)

The formulation of the background currents, *I*_bCa_, *I*_bNa_ and *I*_bCl_, are based on [74] and are given by

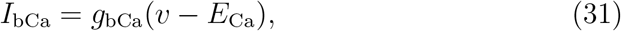

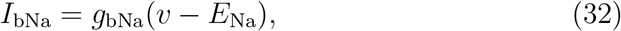

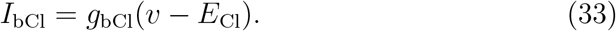

##### Sodium-calcium exchanger current (*I*_NaCa_)

The formulation of the Na^+^−Ca^2+^ exchanger current, *I*_NaCa_, is based on [74] and is given by

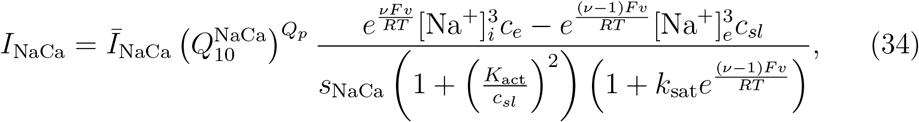

where

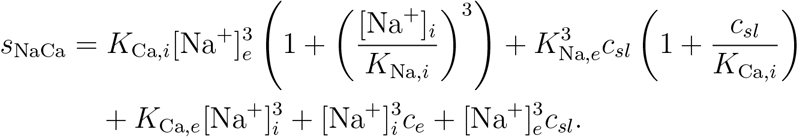

##### Sarcolemmal Ca^2+^ pump current (*I*_pCa_)

The formulation of the current through the sarcolemmal Ca^2+^ pump, *I*_pCa_, is based on [74] and is given by

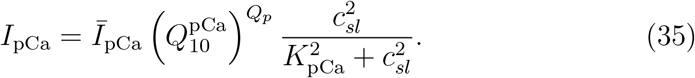

##### Sodium-potassium pump current (*I*_NaK_)

The current through the Na^+^−K^+^ pump, *I*_NaK_, is based on [74] and is given by

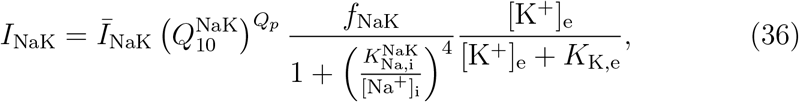

where

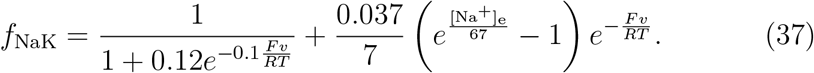

### 6.2 Intracellular [Ca^2+^] dynamics

The Ca^2+^ dynamics are governed by

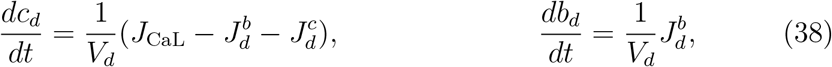

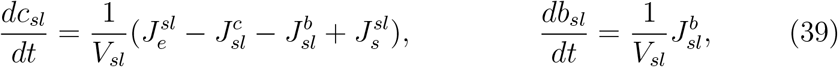

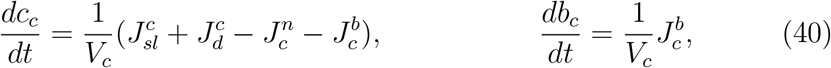

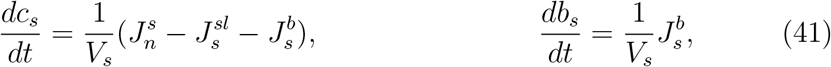

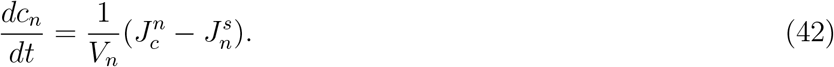

Here, *c_d_* is the concentration of free Ca^2+^ in the dyad, *b_d_* is the concentration of Ca^2+^ bound to a buffer in the dyad, *c_sl_* is the concentration of free Ca^2+^ in the sub-sarcolemmal (SL) compartment, *b_sl_* is the concentration of Ca^2+^ bound to a buffer in the SL compartment, *c_c_* is the concentration of free Ca^2+^ in the bulk cytosol, *b_c_* is the concentration of Ca^2+^ bound to a buffer in the bulk cytosol, *c_s_* is the concentration of free Ca^2+^ in the junctional sarcoplasmic reticulum (jSR), *b_s_* is the concentration of Ca^2+^ bound to a buffer in the jSR, and *c_n_* is the concentration of free Ca^2+^ in the network sarcoplasmic reticulum (nSR). The expressions for the fluxes are specified below.

### 6.3 Ca^2+^ fluxes

#### Flux through the SERCA pumps

The flux from the bulk cytosol to the nSR through the SERCA pumps is based on [74] and given by

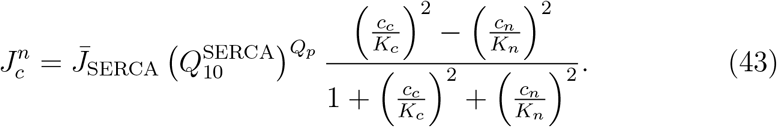

#### Flux through the RyRs

The flux from the jSR to the SL compartment is given by

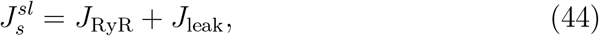

where *J*_RyR_ is the flux through the active RyR channels and *J*_leak_ is the flux through passive RyR channels that are always open, given by

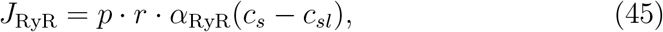

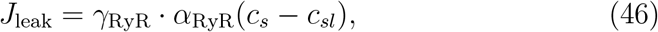

respectively. Here, *p* represents the open probability of the active RyR channels and is given by

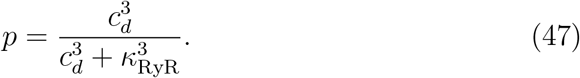

Furthermore, *r* is the fraction of RyR channels that are not inactivated and is governed by the equation

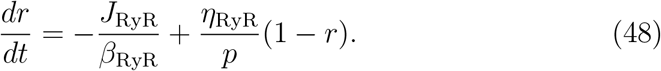

#### Passive diffusion fluxes between compartments

The passive diffusion fluxes between intracellular compartments are given by

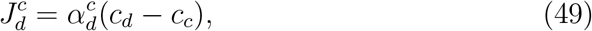

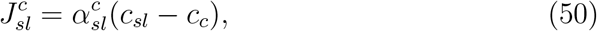

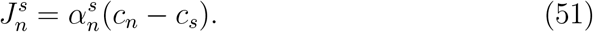

#### Buffer fluxes

The fluxes of free Ca^2+^ binding to a Ca^2+^ buffer are given by

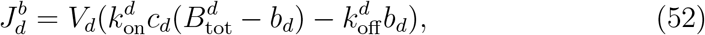

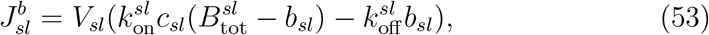

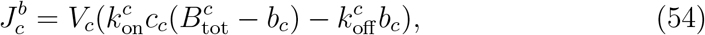

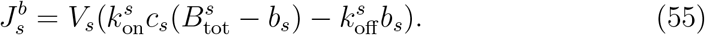

#### Membrane fluxes

The membrane Ca^2+^ fluxes, *J*_CaL_, *J*_bCa_, *J*_pCa_, and *J*_NaCa_, are given by

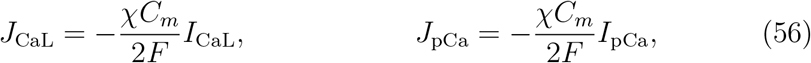

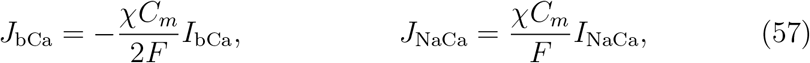

where *I*_CaL_, *I*_bCa_, *I*_pCa_, and *I*_NaCa_ are defined by the expressions given above. Furthermore,

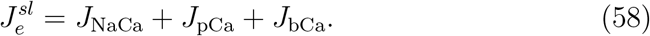

### 6.4 Intracellular Na^+^ dynamics

The intracellular Na^+^ concentration is governed by

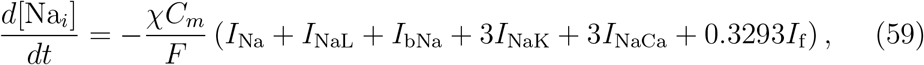

where the currents *I*_Na_, *I*_NaL_, *I*_bNa_, *I*_NaK_, *I*_NaCa_, and *I*_f_ are specified above.

### 6.5 Nernst equilibrium potentials

The Nernst equilibrium potentials for the ion channels are defined as

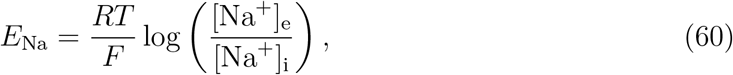

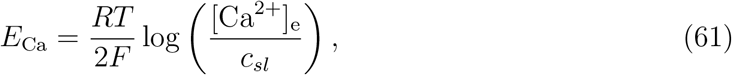

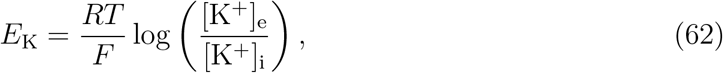

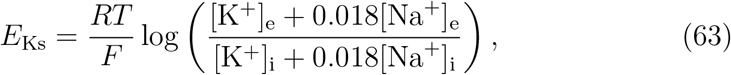

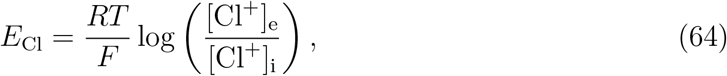

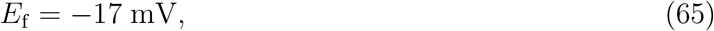

for the parameter values given in Table 8.

### 6.6 Baseline parameter values

### 6.7 Supplementary figures

**Table 7:** Default geometry parameters of the base model.

| Parameter | Description | Value |
| --- | --- | --- |
| $V_d$ | Volume fraction of the dyadic subspace | 0.001 |
| $V_{sl}$ | Volume fraction of the SL compartment | 0.028 |
| $V_c$ | Volume fraction of the bulk cytosol | 0.917 |
| $V_s$ | Volume fraction of the jSR | 0.004 |
| $V_n$ | Volume fraction of the nSR | 0.05 |
| $\chi$ | Cell surface to volume ratio | $0.6 \mu\text{m}^{-1}$ |

**Table 8:** Physical constants and ionic concentrations of the base model.

| Parameter | Description | Value |
| --- | --- | --- |
| $C_m$ | Specific membrane capacitance | $0.01 \text{ pF}/\mu\text{m}^2$ |
| $F$ | Faraday's constant | $96.485 \text{ C}/\text{mmol}$ |
| $R$ | Universal gas constant | $8.314 \text{ J}/(\text{mol}\cdot\text{K})$ |
| $T$ | Temperature | 310 K |
| $[\text{Ca}^{2+}]_e$ | Extracellular $\text{Ca}^{2+}$ concentration | 1.8 mM |
| $[\text{Na}^+]_e$ | Extracellular sodium concentration | 140 mM |
| $[\text{K}^+]_e$ | Extracellular potassium concentration | 5.4 mM |
| $[\text{K}^+]_e^b$ | Baseline extracellular potassium concentration | 5.4 mM |
| $[\text{K}^+]_i$ | Intracellular potassium concentration | 120 mM |
| $[\text{Cl}^-]_e$ | Extracellular chloride concentration | 150 mM |
| $[\text{Cl}^-]_i$ | Intracellular chloride concentration | 15 mM |

**Table 9:** Conductances and similar cell-specific parameter values in the base model formulation. Note that the parameter values of this table define the human left atrial version of the base model. For the human pulmonary vein version, the adjustment factors of Table 10 are applied.

| Parameter | Value | Parameter | Value |
| --- | --- | --- | --- |
| $g_{\text{Na}}$ | 7.056 mS/$\mu\text{F}$ | $\bar{I}_{\text{NaCa}}$ | 2.22 $\mu\text{A}/\mu\text{F}$ |
| $g_{\text{NaL}}$ | 0.021 mS/$\mu\text{F}$ | $\bar{I}_{\text{pCa}}$ | 0.068 $\mu\text{A}/\mu\text{F}$ |
| $g_{\text{to}}$ | 0.55 mS/$\mu\text{F}$ | $\bar{J}_{\text{SERCA}}$ | 0.00024 mM/ms |
| $g_{\text{Kr}}$ | 1.2 mS/$\mu\text{F}$ | $\alpha_{\text{RyR}}$ | 0.0075 ms<sup>-1</sup> |
| $g_{\text{Ks}}$ | 0.21 mS/$\mu\text{F}$ | $\beta_{\text{RyR}}$ | 0.038 mM |
| $g_{\text{K1}}$ | 2.62 mS/$\mu\text{F}$ | $\alpha_d^c$ | 0.0034 ms<sup>-1</sup> |
| $g_{\text{f}}$ | 0.0001 mS/$\mu\text{F}$ | $\alpha_{sl}^c$ | 0.15 ms<sup>-1</sup> |
| $g_{\text{bCl}}$ | 0.0114 mS/$\mu\text{F}$ | $\alpha_n^s$ | 0.012 ms<sup>-1</sup> |
| $g_{\text{CaL}}$ | 0.049 nL/($\mu\text{F}$ ms) | $B_{\text{tot}}^c$ | 0.07 mM |
| $g_{\text{bCa}}$ | 0.000832 mS/$\mu\text{F}$ | $B_{\text{tot}}^d$ | 1.2 mM |
| $g_{\text{bNa}}$ | 0.00185 mS/$\mu\text{F}$ | $B_{\text{tot}}^{sl}$ | 0.9 mM |
| $g_{\text{Kur}}$ | 2.475 mS/$\mu\text{F}$ | $B_{\text{tot}}^s$ | 27 mM |
| $\bar{I}_{\text{NaK}}$ | 2.76 $\mu\text{A}/\mu\text{F}$ | | |

**Table 10:** Adjustment factors for the pulmonary vein (PV) version of the base model, based on [22].

| Parameter | PV scaling factor |
| --- | --- |
| $g_{\text{K1}}$ | 0.58 |
| $g_{\text{Kr}}$ | 1.5 |
| $g_{\text{Ks}}$ | 1.6 |
| $g_{\text{to}}$ | 0.75 |
| $g_{\text{CaL}}$ | 0.7 |

**Table 11:** Parameters for the intracellular Ca^2+^ fluxes of the base model.

| Parameter | Flux | Value |
| --- | --- | --- |
| $K_c$ | $J_c^n$ | 0.00025 mM |
| $K_n$ | $J_c^n$ | 1.7 mM |
| $\gamma_{\text{RyR}}$ | $J_s^{sl}$ | 0.001 |
| $\kappa_{\text{RyR}}$ | $J_{\text{RyR}}$ | 0.015 mM |
| $\eta_{\text{RyR}}$ | $J_s^{sl}$ | 0.00001 ms<sup>-1</sup> |

**Table 12:** Additional parameters for the membrane currents of the base model.

| Parameter | Current | Value |
| --- | --- | --- |
| $k_{\text{sat}}$ | $I_{\text{NaCa}}$ | 0.3 |
| $\nu$ | $I_{\text{NaCa}}$ | 0.3 |
| $K_{\text{act}}$ | $I_{\text{NaCa}}$ | 0.00015 mM |
| $K_{\text{Ca},i}$ | $I_{\text{NaCa}}$ | 0.0036 mM |
| $K_{\text{Ca},e}$ | $I_{\text{NaCa}}$ | 1.3 mM |
| $K_{\text{Na},i}$ | $I_{\text{NaCa}}$ | 12.3 mM |
| $K_{\text{Na},e}$ | $I_{\text{NaCa}}$ | 87.5 mM |
| $K_{\text{Na},i}^{\text{NaK}}$ | $I_{\text{NaK}}$ | (11 mM) · ($Q_{10}^{\text{KNaK}}$)<sup>Q<sub>p</sub></sup> |
| $K_{\text{K},e}$ | $I_{\text{NaK}}$ | 1.5 mM |
| $K_{\text{pCa}}$ | $I_{\text{pCa}}$ | 0.0005 mM |

**Table 13:** Transition rates for the Ca^2+^ buffers of the base model.

| Parameter | Compartment | Value |
| --- | --- | --- |
| $k_{\text{on}}^c$ | Bulk cytosol | 40 ms<sup>-1</sup>mM<sup>-1</sup> |
| $k_{\text{off}}^c$ | Bulk cytosol | 0.03 ms<sup>-1</sup> |
| $k_{\text{on}}^d$ | Dyad | 100 ms<sup>-1</sup>mM<sup>-1</sup> |
| $k_{\text{off}}^d$ | Dyad | 1 ms<sup>-1</sup> |
| $k_{\text{on}}^{sl}$ | Subsarcolemmal space | 100 ms<sup>-1</sup>mM<sup>-1</sup> |
| $k_{\text{off}}^{sl}$ | Subsarcolemmal space | 0.15 ms<sup>-1</sup> |
| $k_{\text{on}}^s$ | Junctional SR | 100 ms<sup>-1</sup>mM<sup>-1</sup> |
| $k_{\text{off}}^s$ | Junctional SR | 65 ms<sup>-1</sup> |

**Table 14:** Transition rates for the *I*_Kr_ Markov model. Here, 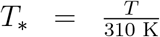 where *T* is the temperature.

|  |  |
| --- | --- |
| $\alpha_{\text{Kr},1}$ | $0.4 \cdot T_* \cdot e^{24.335+T_*^{-1}(0.0112v-25.914)}$ |
| $\beta_{\text{Kr},1}$ | $T_* \cdot e^{13.688+T_*^{-1}(-0.0603v-15.707)}$ |
| $\alpha_{\text{Kr},2}$ | $0.4 \cdot T_* \cdot e^{22.746+T_*^{-1}(-25.914)}$ |
| $\beta_{\text{Kr},2}$ | $T_* \cdot e^{13.193+T_*^{-1}(-15.707)}$ |
| $\alpha_{\text{Kr},3}$ | $0.4 \cdot T_* \cdot e^{22.098+T_*^{-1}(0.0365v-25.914)}$ |
| $\beta_{\text{Kr},3}$ | $T_* \cdot e^{7.313+T_*^{-1}(-0.0399v-15.707)}$ |
| $\alpha_{\text{Kr},4} \text{ (WT)}$ | $T_* \cdot \left( \frac{[\text{K}^+]_e^b}{[\text{K}^+]_e} \right)^{0.4} e^{30.016+T_*^{-1}(0.0223v-30.888)}$ |
| $\alpha_{\text{Kr},4} \text{ (N588K)}$ | $0.25 \cdot T_* \cdot \left( \frac{[\text{K}^+]_e^b}{[\text{K}^+]_e} \right)^{0.4} e^{30.016+T_*^{-1}(0.0223v-30.888)}$ |
| $\beta_{\text{Kr},4} \text{ (WT)}$ | $T_* \cdot e^{30.061+T_*^{-1}(-0.0312v-33.243)}$ |
| $\beta_{\text{Kr},4} \text{ (N588K)}$ | $5 \cdot T_* \cdot e^{30.061+T_*^{-1}(-0.0312v-33.243)}$ |

**Table 15:**
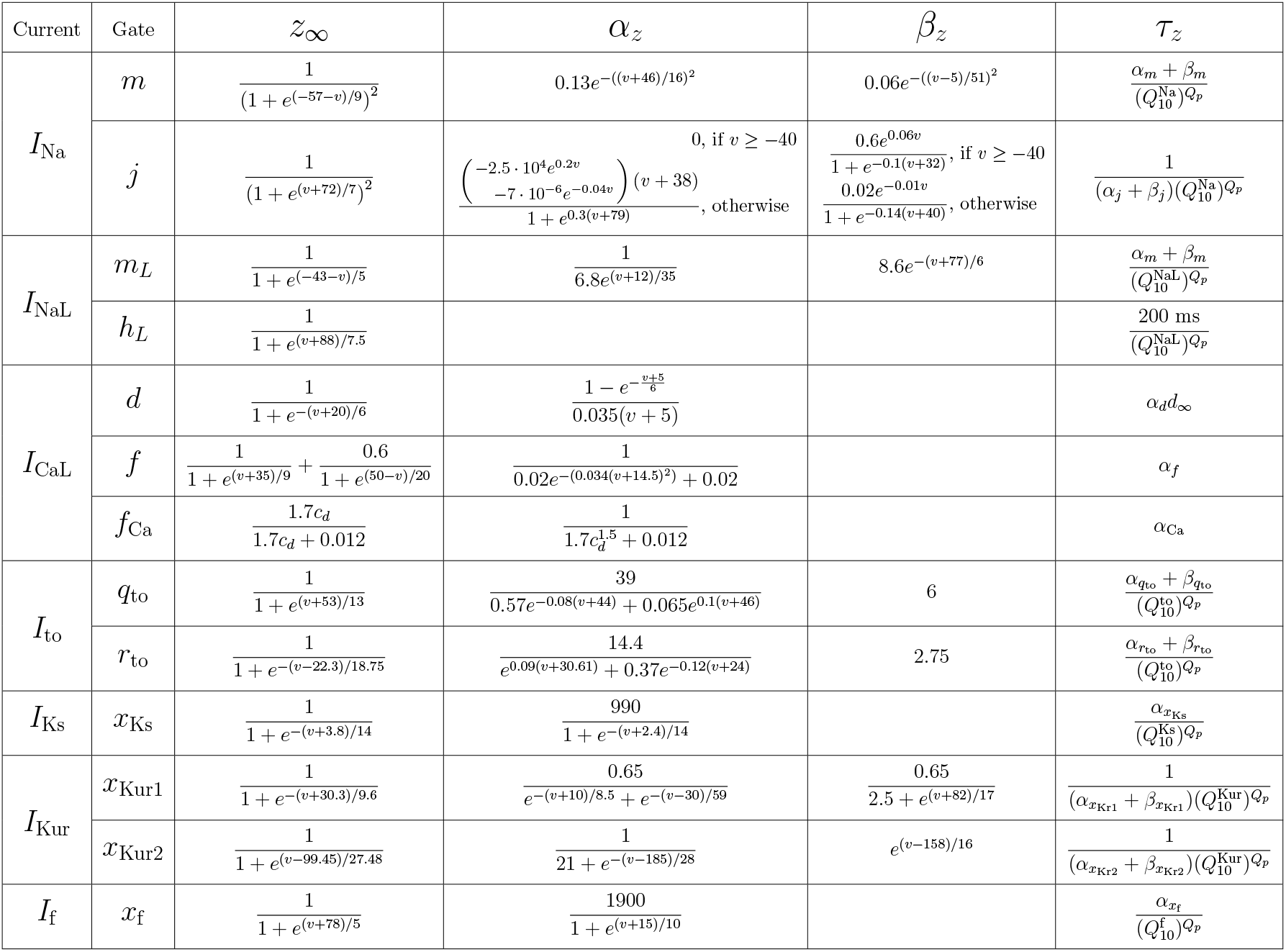
Specification of the parameters *z*_∞_ and *τ_z_*, for *z* = *m, j, m_L_*, *h_L_*, *d, f, f*_Ca_, *q*_to_, *r*_to_, *x*_Ks_, *x*_Kur1_, *x*_Kur2_, and *x*_f_ in the equations for the gating variables (8).

**Figure 9:**
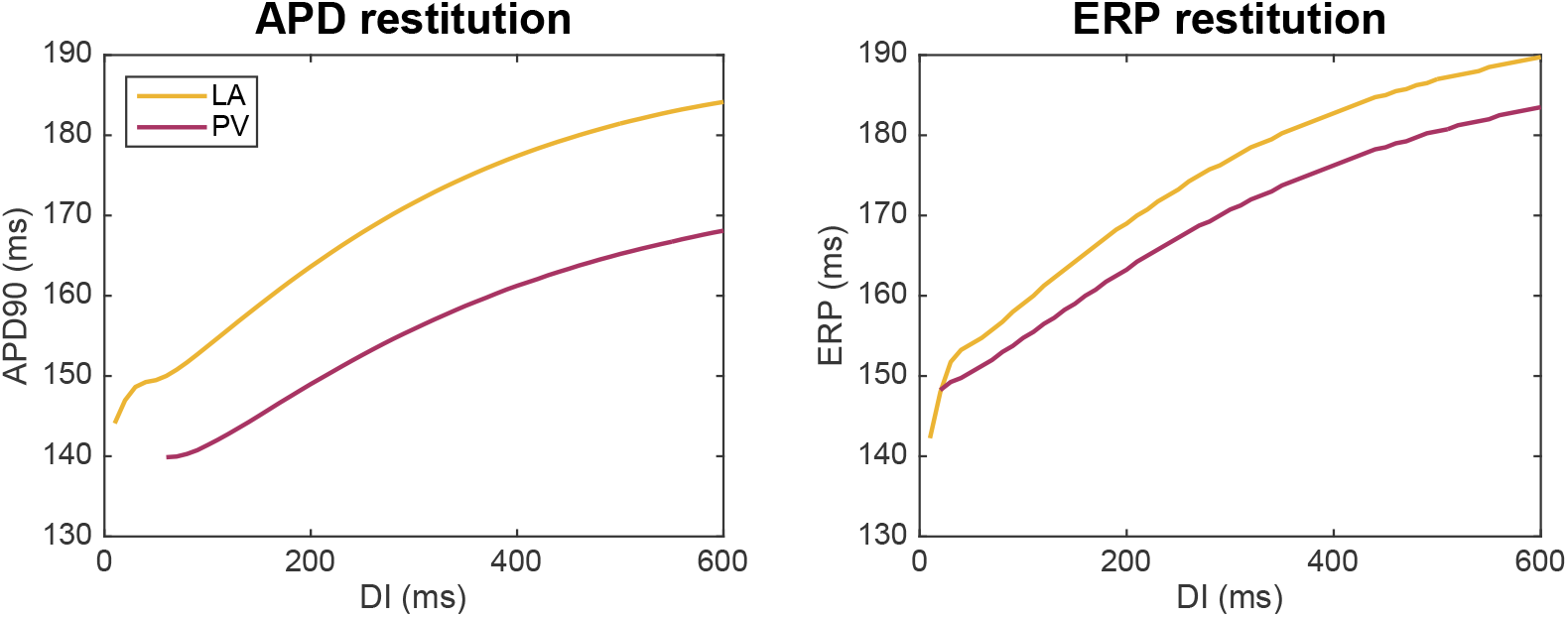
Action potential duration (APD) and effective refractory period (ERP) restitution curves for the LA and PV base models. For the APD restitution curve, the APD90 value is computed for a number of different diastolic interval (DI) values. The diastolic interval is defined as the time between the time point when the membrane potential has returned to 90% repolarization in the previous action potential (resulting from an S1 stimulation) to the time the next stimulation (S2) is applied. To compute the ERP restitution, a third stimulation (S3) is applied for each S2 DI. This S3 stimulation (lasting for 3 ms and of a strength about twice as large as what is needed to generate an action potential from rest) is applied at progressively shorter time intervals after the S2 stimulation and the ERP is defined as the longest S2S3 interval that failed to capture [80].

## Notes

### Competing Interest Statement

The authors have declared no competing interest.

## References

[1] Stanley Nattel and Dobromir Dobrev. Controversies about atrial fibrillation mechanisms: aiming for order in chaos and whether it matters. Circulation Research, 120(9):1396–1398, 2017.

[2] Cristian Martignani, Giulia Massaro, Mauro Biffi, Matteo Ziacchi, and Igor Diemberger. Atrial fibrillation: an arrhythmia that makes healthcare systems tremble. Journal of Medical Economics, 23(7):667–669, 2020. PMID: 32255385.

[3] Julien Feghaly, Patrick Zakka, Barry London, Calum A MacRae, and Marwan M Refaat. Genetics of atrial fibrillation. Journal of the American Heart Association, 7(20):e009884, 2018.

[4] Axel Brandes, Marcelle D Smit, Bao Oanh Nguyen, Michiel Rienstra, and Isabelle C Van Gelder. Risk factor management in atrial fibrillation. Arrhythmia & Electrophysiology Review, 7(2):118, 2018.

[5] Diane Fatkin, Celine F Santiago, Inken G Huttner, Steven A Lubitz, and Patrick T Ellinor. Genetics of atrial fibrillation: state of the art in 2017. Heart, Lung and Circulation, 26(9):894–901, 2017.

[6] Lu-Chen Weng, Sarah R Preis, Olivia L Hulme, Martin G Larson, Seung Hoan Choi, Biqi Wang, Ludovic Trinquart, David D McManus, Laila Staerk, Honghuang Lin, et al. Genetic predisposition, clinical risk factor burden, and lifetime risk of atrial fibrillation. Circulation, 137(10):1027–1038, 2018.

[7] David Calvo, David Filgueiras-Rama, and Jose Jalife. Mechanisms and drug development in atrial fibrillation. Pharmacological Reviews, 70(3):505–525, 2018.

[8] Nicola Jaime Adderley, Ronan Ryan, Krishnarajah Nirantharakumar, and Tom Marshall. Prevalence and treatment of atrial fibrillation in UK general practice from 2000 to 2016. Heart, 105(1):27–33, 2019.

[9] Jussi T Koivumäki, Topi Korhonen, and Pasi Tavi. Impact of sarcoplasmic reticulum calcium release on calcium dynamics and action potential morphology in human atrial myocytes: a computational study. PLoS Computational Biology, 7(1):e1001067, 2011.

[10] Lasse Skibsbye, Thomas Jespersen, Torsten Christ, Mary M Maleckar, Jonas van den Brink, Pasi Tavi, and Jussi T Koivumäki. Refractoriness in human atria: time and voltage dependence of sodium channel availability. Journal of Molecular and Cellular Cardiology, 101:26–34, 2016.

[11] Haibo Ni, Alex Fogli Iseppe, Wayne R Giles, Sanjiv M Narayan, Henggui Zhang, Andrew G Edwards, Stefano Morotti, and Eleonora Grandi. Populations of in silico myocytes and tissues reveal synergy of multiatrialpredominant K^+^-current block in atrial fibrillation. British Journal of Pharmacology, 177(19):4497–4515, 2020.

[12] Caroline H Roney, Jason D Bayer, Hubert Cochet, Marianna Meo, Rémi Dubois, Pierre Jaïs, and Edward J Vigmond. Variability in pulmonary vein electrophysiology and fibrosis determines arrhythmia susceptibility and dynamics. PLoS Computational Biology, 14(5):e1006166, 2018.

[13] Konstantinos N Aronis, Rheeda L Ali, Jialiu A Liang, Shijie Zhou, and Natalia A Trayanova. Understanding AF mechanisms through computational modelling and simulations. Arrhythmia & Electrophysiology Review, 8(3):210, 2019.

[14] Natalia A Trayanova. Mathematical approaches to understanding and imaging atrial fibrillation: significance for mechanisms and management. Circulation Research, 114(9):1516–1531, 2014.

[15] Michael Clerx, Gary R Mirams, Albert J Rogers, Sanjiv M Narayan, and Wayne R Giles. Immediate and delayed response of simulated human atrial myocytes to clinically-relevant hypokalemia. Frontiers in Physiology, 2021.

[16] Oleg V Aslanidi, Michael A Colman, Marta Varela, Jichao Zhao, Bruce H Smaill, Jules C Hancox, Mark R Boyett, and Henggui Zhang. Heterogeneous and anisotropic integrative model of pulmonary veins: computational study of arrhythmogenic substrate for atrial fibrillation. Interface Focus, 3(2):20120069, 2013.

[17] Jordi Heijman, Henry Sutanto, Harry JGM Crijns, Stanley Nattel, and Natalia A Trayanova. Computational models of atrial fibrillation: achievements, challenges, and perspectives for improving clinical care. Cardiovascular Research, 117(7):1682–1699, 2021.

[18] Michel Haissaguerre, Pierre Jaïs, Dipen C Shah, Atsushi Takahashi, Mélèze Hocini, Gilles Quiniou, Stéphane Garrigue, Alain Le Mouroux, Philippe Le Métayer, and Jacques Clémenty. Spontaneous initiation of atrial fibrillation by ectopic beats originating in the pulmonary veins. New England Journal of Medicine, 339(10):659–666, 1998.

[19] Michel Haïssaguerre, Dipen C Shah, Pierre Jaïs, Mélèze Hocini, Teiichi Yamane, Isabel Deisenhofer, Michel Chauvin, Stéphane Garrigue, and Jacques Clémenty. Electrophysiological breakthroughs from the left atrium to the pulmonary veins. Circulation, 102(20):2463–2465, 2000.

[20] Mélèze Hocini, Siew Y Ho, Tokuhiro Kawara, André C Linnenbank, Mark Potse, Dipen Shah, Pierre Jaïs, Michiel J Janse, Michel Haïssaguerre, and Jacques MT De Bakker. Electrical conduction in canine pulmonary veins: electrophysiological and anatomic correlation. Circulation, 105(20):2442–2448, 2002.

[21] Antony J Workman, Kathleen A Kane, and Andrew C Rankin. Cellular bases for human atrial fibrillation. Heart Rhythm, 5(6):S1–S6, 2008.

[22] Joachim R Ehrlich, Tae-Joon Cha, Liming Zhang, Denis Chartier, Peter Melnyk, Stefan H Hohnloser, and Stanley Nattel. Cellular electrophysiology of canine pulmonary vein cardiomyocytes: action potential and ionic current properties. The Journal of Physiology, 551(3):801–813, 2003.

[23] Marta Varela, Michael A Colman, Jules C Hancox, and Oleg V Aslanidi. Atrial heterogeneity generates re-entrant substrate during atrial fibrillation and anti-arrhythmic drug action: mechanistic insights from canine atrial models. PLoS Computational Biology, 12(12):e1005245, 2016.

[24] Justus M Anumonwo and Sandeep V Pandit. Ionic mechanisms of arrhythmogenesis. Trends in Cardiovascular Medicine, 25(6):487–496, 2015.

[25] Vincent Jacquemet and Craig S Henriquez. Genesis of complex fractionated atrial electrograms in zones of slow conduction: a computer model of microfibrosis. Heart Rhythm, 6(6):803–810, 2009.

[26] Piero C Franzone, Luca F Pavarino, and Simone Scacchi. Mathematical Cardiac Electrophysiology, volume 13. Springer, 2014.

[27] Olaf Dössel, Martin W Krueger, Frank M Weber, Mathias Wilhelms, and Gunnar Seemann. Computational modeling of the human atrial anatomy and electrophysiology. Medical & Biological Engineering & Computing, 50(8):773–799, 2012.

[28] Aslak Tveito, Karoline H Jæger, Miroslav Kuchta, Kent-Andre Mardal, and Marie E Rognes. A cell-based framework for numerical modeling of electrical conduction in cardiac tissue. Frontiers in Physics, 5:48, 2017.

[29] Karoline Horgmo Jæger, Andrew G Edwards, Andrew McCulloch, and Aslak Tveito. Properties of cardiac conduction in a cell-based computational model. PLoS Computational Biology, 15(5):e1007042, 2019.

[30] Karoline Horgmo Jæger, Kristian Gregorius Hustad, Xing Cai, and Aslak Tveito. Efficient numerical solution of the EMI model representing the extracellular space (E), cell membrane (M) and intracellular space (I) of a collection of cardiac cells. Frontiers in Physics, 8:539, 2021.

[31] Karoline Horgmo Jæger and Aslak Tveito. Derivation of a cell-based mathematical model of excitable cells. In Modeling Excitable Tissue, pages 1–13. Springer, Cham, 2020.

[32] Karoline Horgmo Jæger, Andrew G. Edwards, Wayne R. Giles, and Aslak Tveito. From millimeters to micrometers; re-introducing myocytes in models of cardiac electrophysiology. Preprint, 2021.

[33] Madison S Spach, J Francis Heidlage, Paul C Dolber, and Roger C Barr. Mechanism of origin of conduction disturbances in aging human atrial bundles: experimental and model study. Heart Rhythm, 4(2):175–185, 2007.

[34] Steven Niederer, Lawrence Mitchell, Nicolas Smith, and Gernot Plank. Simulating human cardiac electrophysiology on clinical time-scales. Frontiers in Physiology, 2:14, 2011.

[35] Steven A Niederer, Eric Kerfoot, Alan P Benson, Miguel O Bernabeu, Olivier Bernus, Chris Bradley, Elizabeth M Cherry, Richard Clayton, Flavio H Fenton, Alan Garny, et al. Verification of cardiac tissue electrophysiology simulators using an n-version benchmark. Philosophical Transactions of the Royal Society A: Mathematical, Physical and Engineering Sciences, 369(1954):4331–4351, 2011.

[36] RH Clayton and AV Panfilov. A guide to modelling cardiac electrical activity in anatomically detailed ventricles. Progress in Biophysics and Molecular Biology, 96(1-3):19–43, 2008.

[37] Fagen Xie, Zhilin Qu, Junzhong Yang, Ali Baher, James N Weiss, Alan Garfinkel, et al. A simulation study of the effects of cardiac anatomy in ventricular fibrillation. The Journal of Clinical Investigation, 113(5):686–693, 2004.

[38] Mark J McPate, Rona S Duncan, James T Milnes, Harry J Witchel, and Jules C Hancox. The N588K-HERG K^+^ channel mutation in the ‘short QT syndrome’: mechanism of gain-in-function determined at 37 °C. Biochemical and Biophysical Research Communications, 334(2):441–449, 2005.

[39] Karoline Horgmo Jæger, Samuel Wall, and Aslak Tveito. Computational prediction of drug response in short QT syndrome type 1 based on measurements of compound effect in stem cell-derived cardiomyocytes. PLoS Computational Biology, 17(2):e1008089, 2021.

[40] KUI Hong, Preben Bjerregaard, Ihor Gussak, and Ramon Brugada. Short QT syndrome and atrial fibrillation caused by mutation in KCNH2. Journal of Cardiovascular Electrophysiology, 16(4):394–396, 2005.

[41] Morten Salling Olesen, Lena Refsgaard, Anders Gaarsdal Holst, Anders Peter Larsen, Søren Grubb, Stig Haunsø, Jesper Hastrup Svendsen, Søren-Peter Olesen, Nicole Schmitt, and Kirstine Calloe. A novel KCND3 gain-of-function mutation associated with early-onset of persistent lone atrial fibrillation. Cardiovascular Research, 98(3):488–495, 2013.

[42] Makarand Deo, Yanfei Ruan, Sandeep V Pandit, Kushal Shah, Omer Berenfeld, Andrew Blaufox, Marina Cerrone, Sami F Noujaim, Marco Denegri, José Jalife, and Silvia G Priori. KCNJ2 mutation in short QT syndrome 3 results in atrial fibrillation and ventricular proarrhythmia. Proceedings of the National Academy of Sciences, 110(11):4291–4296, 2013.

[43] Timothy M Olson, Alexey E Alekseev, Xiaoke K Liu, Sungjo Park, Leonid V Zingman, Martin Bienengraeber, Srinivasan Sattiraju, Jeffrey D Ballew, Arshad Jahangir, and Andre Terzic. Kv1.5 channelopathy due to KCNA5 loss-of-function mutation causes human atrial fibrillation. Human Molecular Genetics, 15(14):2185–2191, 2006.

[44] Pengyun Wang, Qinbo Yang, Xiaofen Wu, Yanzong Yang, Lisong Shi, Chuchu Wang, Gang Wu, Yunlong Xia, Bo Yang, Rongfeng Zhang, et al. Functional dominant-negative mutation of sodium channel subunit gene SCN3B associated with atrial fibrillation in a chinese GeneID population. Biochemical and Biophysical Research Communications, 398(1):98–104, 2010.

[45] Isabelle L Thibodeau, Ji Xu, Qiuju Li, Gele Liu, Khanh Lam, John P Veinot, David H Birnie, Douglas L Jones, Andrew D Krahn, Robert Lemery, Bruce J Nicholson, and Michael H Gollob. Paradigm of genetic mosaicism and lone atrial fibrillation: physiological characterization of a connexin 43-deletion mutant identified from atrial tissue. Circulation, 122(3):236–244, 2010.

[46] Karoline Horgmo Jæger, Verena Charwat, Bérénice Charrez, Henrik Finsberg, Mary M Maleckar, Sam Wall, Kevin Healy, and Aslak Tveito. Improved computational identification of drug response using optical measurements of human stem cell derived cardiomyocytes in microphysiological systems. Frontiers in Pharmacology, 10:1648, 2020.

[47] Aslak Tveito, Karoline Horgmo Jæger, Mary M Maleckar, Wayne R Giles, and Samuel Wall. Computational translation of drug effects from animal experiments to human ventricular myocytes. Scientific Reports, 10(1):1–11, 2020.

[48] Karoline Horgmo Jæger, Andrew G. Edwards, Wayne R. Giles, and Aslak Tveito. A computational method for identifying an optimal combination of existing drugs to repair the action potentials of SQT1 ventricular myocytes. PLoS Computational Biology, 17(8):e1009233, 2021.

[49] Fiorenzo Gaita, Carla Giustetto, Francesca Bianchi, Christian Wolpert, Rainer Schimpf, Riccardo Riccardi, Stefano Grossi, Elena Richiardi, and Martin Borggrefe. Short QT syndrome: a familial cause of sudden death. Circulation, 108(8):965–970, 2003.

[50] Martin Fink, Denis Noble, Laszlo Virag, Andras Varro, and Wayne R Giles. Contributions of HERG K^+^ current to repolarization of the human ventricular action potential. Progress in Biophysics and Molecular Biology, 96(1-3):357–376, 2008.

[51] Eleonora Grandi, Sandeep V Pandit, Niels Voigt, Antony J Workman, Dobromir Dobrev, José Jalife, and Donald M Bers. Human atrial action potential and Ca^2+^ model: sinus rhythm and chronic atrial fibrillation. Circulation Research, 109(9):1055–1066, 2011.

[52] Jeroen G Stinstra, Sarah F Roberts, John B Pormann, Rob S MacLeod, and Craig S Henriquez. A model of 3D propagation in discrete cardiac tissue. In Computers in Cardiology, 2006, pages 41–44. IEEE, 2006.

[53] Jeroen Stinstra, Rob MacLeod, and Craig Henriquez. Incorporating histology into a 3D microscopic computer model of myocardium to study propagation at a cellular level. Annals of Biomedical Engineering, 38(4):1399–1414, 2010.

[54] R. Anderson, J. Andrej, A. Barker, J. Bramwell, J.-S. Camier, J. Cerveny V. Dobrev, Y. Dudouit, A. Fisher, Tz. Kolev, W. Pazner, M. Stowell, V. Tomov, I. Akkerman, J. Dahm, D. Medina, and S. Zampini. MFEM: A modular finite element library. Computers & Mathematics with Applications, 2020.

[55] MFEM: Modular finite element methods [Software]. mfem.org.

[56] Karoline Horgmo Jæger, Kristian Gregorius Hustad, Xing Cai, and Aslak Tveito. Operator splitting and finite difference schemes for solving the EMI model. In Modeling Excitable Tissue, pages 44–55. Springer, Cham, 2020.

[57] Teiichi Yamane, Dipen C Shah, Pierre Jaïs, Mélèze Hocini, Jing Tian Peng, Isabel Deisenhofer, Jacques Clémenty, and Michel Haïssaguerre. Dilatation as a marker of pulmonary veins initiating atrial fibrillation. Journal of Interventional Cardiac Electrophysiology, 6(3):245–249, 2002.

[58] Anders Nygren, Céline Fiset, Ludwik Firek, John W Clark, Douglas S Lindblad, Robert B Clark, and Wayne R Giles. Mathematical model of an adult human atrial cell: the role of K^+^ currents in repolarization. Circulation Research, 82(1):63–81, 1998.

[59] RM Ludatscher. Fine structure of the muscular wall of rat pulmonary veins. Journal of Anatomy, 103(Pt 2):345, 1968.

[60] SY Ho, JA Cabrera, VH Tran, J Farre, RH Anderson, and D Sanchez-Quintana. Architecture of the pulmonary veins: relevance to radiofrequency ablation. Heart, 86(3):265–270, 2001.

[61] Siew Yen Ho, Damian Sanchez-Quintana, Jose Angel Cabrera, and Robert H Anderson. Anatomy of the left atrium: implications for radiofrequency ablation of atrial fibrillation. Journal of Cardiovascular Electrophysiology, 10(11):1525–1533, 1999.

[62] Akira Hamabe, Yuji Okuyama, Yasushi Miyauchi, Shengmei Zhou, Hui-Nam Pak, Hrayr S Karagueuzian, Michael C Fishbein, and Peng-Sheng Chen. Correlation between anatomy and electrical activation in canine pulmonary veins. Circulation, 107(11):1550–1555, 2003.

[63] Michael A Colman, Oleg V Aslanidi, Sanjay Kharche, Mark R Boyett, Clifford Garratt, Jules C Hancox, and Henggui Zhang. Pro-arrhythmogenic effects of atrial fibrillation-induced electrical remodelling: insights from the three-dimensional virtual human atria. The Journal of Physiology, 591(17):4249–4272, 2013.

[64] Eugene Patterson, Sunny S Po, Benjamin J Scherlag, and Ralph Lazzara. Triggered firing in pulmonary veins initiated by in vitro autonomic nerve stimulation. Heart Rhythm, 2(6):624–631, 2005.

[65] Eugene Patterson, Ralph Lazzara, Bela Szabo, Hong Liu, David Tang, Yu-Hua Li, Benjamin J Scherlag, and Sunny S Po. Sodium-calcium exchange initiated by the ca2+ transient: an arrhythmia trigger within pulmonary veins. Journal of the American College of Cardiology, 47(6):1196–1206, 2006.

[66] Alejandro Perez-Lugones, James T McMahon, Norman B Ratliff, Walid I Saliba, Robert A Schweikert, Nassir F Marrouche, Eduardo B Saad, José L Navia, Patrick M McCarthy, Patrick Tchou, et al. Evidence of specialized conduction cells in human pulmonary veins of patients with atrial fibrillation. Journal of Cardiovascular Electrophysiology, 14(8):803–809, 2003.

[67] Elizabeth M Cherry, Joachim R Ehrlich, Stanley Nattel, and Flavio H Fenton. Pulmonary vein reentry—properties and size matter: insights from a computational analysis. Heart Rhythm, 4(12):1553–1562, 2007.

[68] Yunfan Gong, Fagen Xie, Kenneth M Stein, Alan Garfinkel, Calin A Culianu, Bruce B Lerman, and David J Christini. Mechanism underlying initiation of paroxysmal atrial flutter/atrial fibrillation by ectopic foci: a simulation study. Circulation, 115(16):2094–2102, 2007.

[69] Minki Hwang, Byounghyun Lim, Jun-Seop Song, Hee Tae Yu, Ah-Jin Ryu, Young-Seon Lee, Boyoung Joung, Eun Bo Shim, and Hui-Nam Pak. Ganglionated plexi stimulation induces pulmonary vein triggers and promotes atrial arrhythmogenecity: In silico modeling study. PLoS One, 12(2):e0172931, 2017.

[70] Michael A Colman, Marta Varela, Jules C Hancox, Henggui Zhang, and Oleg V Aslanidi. Evolution and pharmacological modulation of the arrhythmogenic wave dynamics in canine pulmonary vein model. Europace, 16(3):416–423, 2014.

[71] Oleg V Aslanidi, Michael A Colman, Jonathan Stott, Halina Dobrzynski, Mark R Boyett, Arun V Holden, and Henggui Zhang. 3D virtual human atria: A computational platform for studying clinical atrial fibrillation. Progress in Biophysics and Molecular Biology, 107(1):156–168, 2011.

[72] Julie K Shade, Rheeda L Ali, Dante Basile, Dan Popescu, Tauseef Akhtar, Joseph E Marine, David D Spragg, Hugh Calkins, and Natalia A Trayanova. Preprocedure application of machine learning and mechanistic simulations predicts likelihood of paroxysmal atrial fibrillation recurrence following pulmonary vein isolation. Circulation: Arrhythmia and Electrophysiology, 13(7):e008213, 2020.

[73] Stavros Stavrakis and Sunny Po. Ganglionated plexi ablation: physiology and clinical applications. Arrhythmia & electrophysiology review, 6(4):186, 2017.

[74] Eleonora Grandi, Francesco S Pasqualini, and Donald M Bers. A novel computational model of the human ventricular action potential and Ca transient. Journal of Molecular and Cellular Cardiology, 48(1):112–121, 2010.

[75] Thomas O’Hara, László Virág, András Varró, and Yoram Rudy. Simulation of the undiseased human cardiac ventricular action potential: Model formulation and experimental validation. PLoS Computational Biology, 7(5):e1002061, 2011.

[76] Michelangelo Paci, Jari Hyttinen, Katriina Aalto-Setälä, and Stefano Severi. Computational models of ventricular-and atrial-like human induced pluripotent stem cell derived cardiomyocytes. Annals of Biomedical Engineering, 41(11):2334–2348, 2013.

[77] Marc Courtemanche, Rafael J Ramirez, and Stanley Nattel. Ionic mechanisms underlying human atrial action potential properties: insights from a mathematical model. American Journal of Physiology-Heart and Circulatory Physiology, 275(1):H301–H321, 1998.

[78] Ingrid E Christophersen, Morten S Olesen, Bo Liang, Martin N Andersen, Anders P Larsen, Jonas B Nielsen, Stig Haunsø, Søren-Peter Olesen, Arnljot Tveit, Jesper H Svendsen, and Nicole Schmitt. Genetic variation in KCNA5: impact on the atrial-specific potassium current I_Kur_ in patients with lone atrial fibrillation. European Heart Journal, 34(20):1517–1525, 2013.

[79] Mary M Maleckar, Joseph L Greenstein, Natalia A Trayanova, and Wayne R Giles. Mathematical simulations of ligand-gated and cell-type specific effects on the action potential of human atrium. Progress in Biophysics and Molecular Biology, 98(2-3):161–170, 2008.

[80] Fagen Xie, Zhilin Qu, Alan Garfinkel, and James N Weiss. Electrical refractory period restitution and spiral wave reentry in simulated cardiac tissue. American Journal of Physiology-Heart and Circulatory Physiology, 283(1):H448–H460, 2002.

